# The cryo-EM structure of the CENP-A nucleosome in complex with ggKNL2

**DOI:** 10.1101/2022.06.24.497480

**Authors:** Honghui Jiang, Mariko Ariyoshi, Reito Watanabe, Fumiaki Makino, Keiichi Namba, Tatsuo Fukagawa

**Author notes:** Correspondence to: Tatsuo Fukagawa Graduate School of Frontier Biosciences, Osaka University, Suita, Osaka 565-0871, Japan. These authors equally contribute to this work.

## Abstract

Centromere protein A (CENP-A) nucleosome is an epigenetic marker that specifies centromere position. The Mis18 complex is a licensing factor for new CENP-A deposition via the CENP-A chaperone, Holliday junction recognition protein (HJURP) on the centromere chromatin. Chicken KINETOCHORE NULL2 (KNL2) (ggKNL2), a Mis18 complex component, has a CENP-C-like motif, and our previous study suggested that ggKNL2 directly binds to the CENP-A nucleosome to recruit HJURP/CENP-A to the centromere. However, the molecular basis for CENP-A nucleosome recognition by ggKNK2 remains unclear. Here, we present the cryo-EM structure of the chicken CENP-A nucleosome in complex with a ggKNL2 fragment containing a CENP-C-like motif. Chicken KNL2 distinguishes between CENP-A and histone H3 in the nucleosome using the CENP-C-like motif and its downstream region. Both the C-terminal tail and RG-loop of CENP-A are simultaneously recognized as CENP-A characteristics. The CENP-A nucleosome-ggKNL2 interaction is thus essential for CENP-A deposition. Furthermore, our structural, biochemical, and cell biology data indicate that ggKNL2 alters its binding partner at the centromere during chicken cell cycle progression.

## Introduction

In most organisms, the position of the centromere is defined through sequence-independent epigenetic mechanisms. The histone H3 variant centromere protein A (CENP-A) is a key epigenetic marker of the centromere specification and maintenance (Allshire & Karpen, 2008; Black & Cleveland, 2011; Fukagawa & Earnshaw, 2014; Perpelescu & Fukagawa, 2011; Westhorpe & Straight, 2013). CENP-A forms a nucleosome with other canonical histones (H2A/B and H4) (Black & Cleveland, 2011; Tachiwana *et al*, 2011) and downstream kinetochore components associated with the CENP-A nucleosome to form a functional kinetochore (Amano *et al*, 2009; Foltz *et al*, 2006; Hori *et al*, 2008; Izuta *et al*, 2006; Nishino *et al*, 2012; Okada *et al*, 2006; Weir *et al*, 2016; Yan *et al*, 2019). Therefore, it is crucial to understand the mechanisms by which CENP-A is deposited on the centromeres for their maintenance.

Before CENP-A deposition onto the centromeres, soluble CENP-A-histone H4 dimers specifically bind to Holliday junction recognition protein (HJURP), a CENP-A-specific chaperone, in vertebrate cells (Dunleavy *et al*, 2009; Foltz *et al*, 2009). As CENP-A deposition is not coupled with DNA replication, new CENP-A deposition occurs during the G1 phase in vertebrate cells (Jansen *et al*, 2007). For CENP-A deposition, centromere chromatin must be licensed prior to the G1 phase. The Mis18 complex, containing Mis18α, Mis18β, and M18BP1/KINETOCHORE NULL2 (KNL2), is a licensing factor for CENP-A deposition in vertebrate cells (Fujita *et al*, 2007). The Mis18 gene was originally identified in the fission yeast *Mis18* mutant, which exhibits defects in CENP-A deposition (Hayashi *et al*, 2004). The two human homologues of fission yeast Mis18 were identified as Mis18α and Mis18β. Furthermore, biochemical screening for Mis18 binding proteins identified M18BP1 (Fujita *et al*., 2007), which shows sequence homology to *C. elegans* centromere protein KNL2 (Maddox *et al*, 2007). CENP-A chaperon HJURP directly binds to the Mis18α-Mis18β heterotetramer, which ensures correct CENP-A deposition onto the centromeres (Nardi *et al*, 2016; Subramanian *et al*, 2016). Based on these data, the Mis18 complex is believed to localize to the centromeric region to ensure the deposition of new CENP-A (Pan *et al*, 2019).

Localization of the Mis18 complex to the centromere depends on cell cycle progression in some organisms. Interestingly, this localization is positively regulated by polo-like kinase 1 (PLK1) (McKinley & Cheeseman, 2014) and negatively regulated by cyclin-dependent kinase (CDK) in human cells (Pan *et al*, 2017; Silva *et al*, 2012) to ensure the timely deposition of CENP-A. M18BP1/KNL2 is a component of the Mis18 complex and binds to centromere protein C (CENP-C) in human and mouse cells (Dambacher *et al*, 2012; Moree *et al*, 2011). A recent model suggests that for new CENP-A deposition in human cells, KNL2 localizes to centromeric chromatin via CENP-C binding (McKinley & Cheeseman, 2014; Nardi *et al*., 2016). However, the depletion of CENP-C in *Xenopus* egg extract or chicken DT40 cells has no effect on the association of KNL2 with interphase centromeres (Perpelescu *et al*, 2015; Westhorpe *et al*, 2015). In addition, recent studies have revealed that KNL2 in various species, except humans and mice, has a CENP-C-like motif (French & Straight, 2019; French *et al*, 2017; Hori *et al*, 2017; Kral, 2015; Sandmann *et al*, 2017), which is similar to the conserved CENP-A binding motif of CENP-C (CENP-C motif) (Ariyoshi *et al*, 2021; Kato *et al*, 2013a; Watanabe *et al*, 2019). These data suggest that KNL2 associates with centromeric chromatin by directly binding to the CENP-A nucleosome in these species. In fact, the KNL2 of chicken and *Xenopus* directly binds to the CENP-A nucleosome in vitro (French & Straight, 2019; French *et al*., 2017; Hori *et al*., 2017). Thus, the binding of KNL2 to the CNEP-A nucleosome implies an alternative mechanism of targeting the Mis18 complex for new CENP-A deposition. However, it remains unclear whether and how KNL2 associates with CENP-A nucleosomes in the presence of CENP-C, which also directly interacts with CENP-A. We recently found that the CENP-C motif of CENP-C did not interact with CENP-A in chicken interphase cells, but was directly bound to the CENP-A nucleosome during mitosis upon phosphorylation of the CENP-C near the motif (Ariyoshi *et al*., 2021; Watanabe *et al*., 2019). Therefore, KNL2 was presumably allowed to occupy the CENP-A nucleosome instead of CENP-C in the G1 phase for new CENP-A deposition in chicken cells. However, the detailed molecular mechanism of how chicken KNL2 (ggKNL2) recognizes the CENP-A nucleosome and localizes to the centromere in the presence of CENP-C or other centromeric proteins is unknown.

To address this question, we performed biochemical and structural analyses on the chicken CENP-A nucleosome complexed with the ggKNL2 fragment containing the CENP-C-like motif. In this study, we first confirmed that the CENP-C-like motif of ggKNL2 specifically binds to the CENP-A nucleosome, but not to the H3 nucleosome. We determined the three-dimensional structure of the CENP-A nucleosome complexed with the ggKNL2 fragment containing the CENP-C-like motif at 3.42 Å resolution using cryo-electron microscopy (cryo-EM) single particle image analysis. Our cryo-EM structure demonstrated that ggKNL2 clearly distinguishes CENP-A from histone H3 in nucleosomes. The CENP-C-like motif of ggKNL2 interacts with an acidic patch of the H2A/B and the C-terminal tail of CENP-A in the CENP-A nucleosome. More importantly, ggKNL2 makes a contact with the RG-loop of the CENP-A nucleosome, which is present only in the CENP-A nucleosome but not in the H3 nucleosome. The RG-loop binding of ggKNL2 is suggested to contribute to its specificity toward the CENP-A nucleosome and was reported to be conserved between the CENP-C-like motif containing KNL2 and CNEP-C. ggKNL2 mutants lacking binding to either H2A/B, CENP-A C-terminal tail, or RG loop, does not properly localize in interphase centromeres. In addition, the expression of these mutants did not suppress cell death in KNL2-deficient DT40 cells. This suggests that the binding of KNL2 to the CENP-A nucleosome is essential for the deposition of new CENP-A, which is critical for cell viability. Furthermore, we found that the direct interaction between ggKNL2 and the CENP-A nucleosome is essential for centromere localization in interphase, whereas ggKNL2 localization to mitotic kinetochores depended on CENP-C in DT40 cells. Combined with biochemical and cell biology data, the cryo-EM structure of the ggKNL2-CENP-A nucleosome complex provides new insights into the formation of centromere-specific chromatin.

## Results

### ggKNL2 stably binds to the CENP-A nucleosome via the CENP-C-like motif

ggKNL2 has multiple domains, including SANT and SANTA domains, in addition to the CENP-C-like motif (Fig. 1A) (French *et al*., 2017; Hori *et al*., 2017; Kral, 2015; Sandmann *et al*., 2017). We have previously shown that the CENP-C-like motif of ggKNL2 (amino acid residues 523-543) is critical for centromere localization and binding to the CENP-A nucleosome (Hori *et al*., 2017). To further understand the CENP-A-recognition mechanism of ggKNL2, we assessed the CENP-A-binding ability of ggKNL2, focusing on the regions around the CENP-C-like motif. Various ggKNL2 constructs, with or without the CENP-C-like motif, were prepared as GST-fused recombinant proteins for biochemical assays (Fig. 1A). For the in vitro reconstitution of the CENP-A nucleosome, we used a chimeric CENP-A, in which the N-terminal region of the chicken CENP-A was replaced with the histone H3 N-terminal region. The chimeric CENP-A was previously shown to be replaceable with the wild-type CENP-A in DT40 cells and to form a nucleosome in vitro (Hori *et al*, 2020: Ariyoshi, 2021 #5749; Watanabe *et al*., 2019).

**Figure 1.**
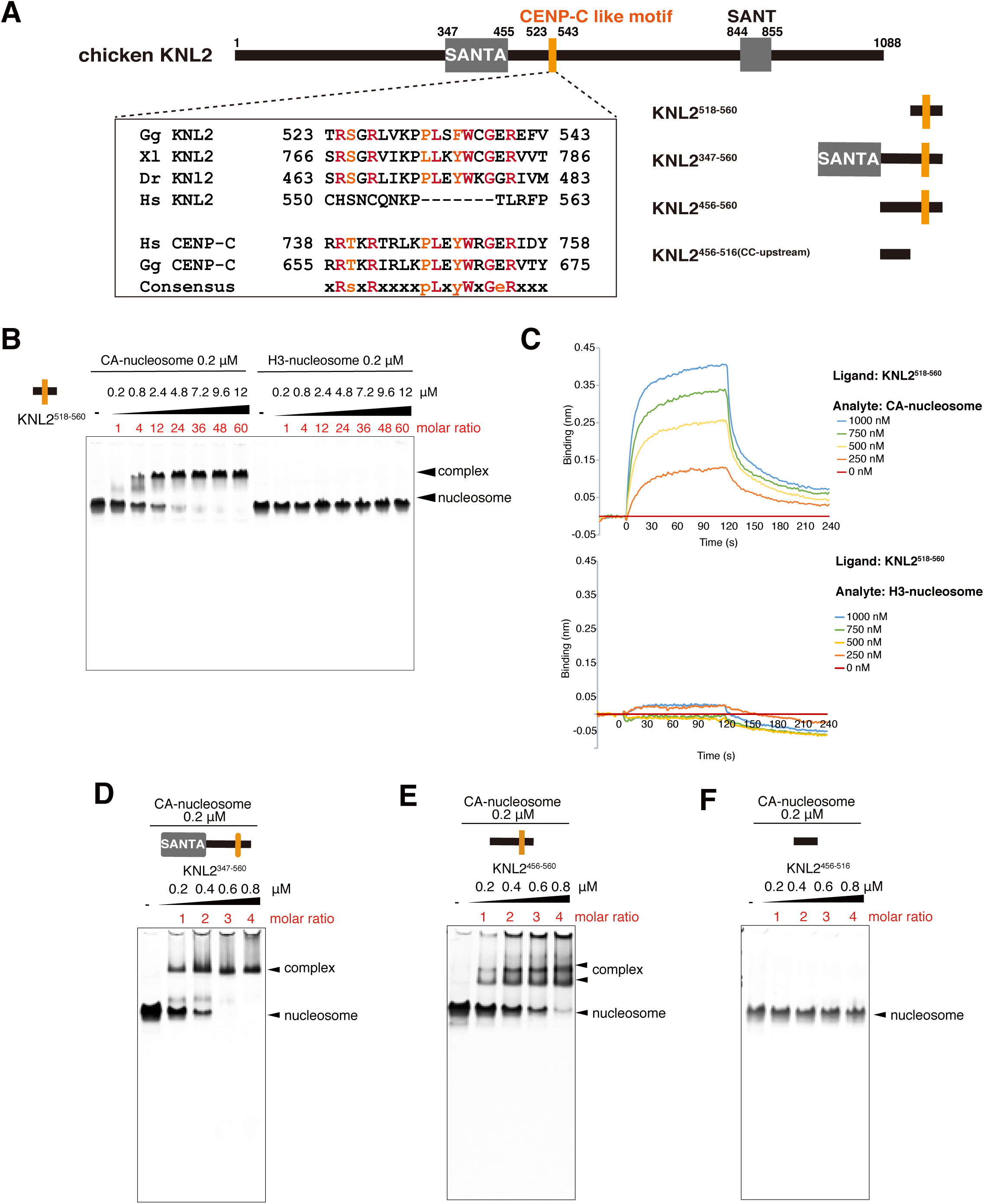
ggKNL2 recognizes the CENP-A nucleosome via the CENP-C like motif A) Schematic diagram of the domain organization of chicken KNL2 (ggKNL2). The CENP-C like motif for CENP-A binding (aa 523-543) is colored in orange. Sequence alignment of the CNEP-C motif of KNL2 or CENP-C from various species: Gg, chicken; Xl, frog; Dr, zebrafish; and Hs, human. The identical and similar residues are colored in red and orange, respectively. The KNL2 fragments, which were used for in vitro CENP-A nucleosome binding assays, are diagramed. **B)** Electrophoretic mobility shift assay (EMSA) to examine binding affinities of the GST-KNL2^518-560^ fragment for the CENP-A or H3 nucleosome. The molar ratio of KNL2^518-560^ against nucleosomes are indicated. **C)** Biolayer interferometry (BLI) assay for the interaction of KNL2^518-560^ with the CENP-A nucleosome (upper panel) or the H3 nucleosome (lower panel). The representative BLI sensorgrams for an analyte (CENP-A or H3 nucleosome) concentration range from 0 to 1 μM are plotted. **D)-F)** EMSA to examine binding affinities of the GST-KNL2 fragment for the CENP-A nucleosome: D, GST-KNL2^347-560^; E, GST-KNL2^456-560^; F, GST-KNL2^456-516^ ^(CC-upstream)^. See also Fig. EV1.

Using electrophoretic mobility shift assays (EMSAs) and biolayer interferometry (BLI) assays, we examined the binding of the various KNL2 fragments (Fig. 1A): KNL2^518-560^ containing the CENP-C-like motif only; KNL2^347-560^ containing the SANTA and CENP-C-like motif; KNL2^456-560^ containing the CENP-C-like motif and its N-terminal upstream (CC- upstream) region but not the SANTA domain, to the CENP-A nucleosome. CENP-A binding assays were also performed using KNL2^456-516^ ^(CC-upstream)^ containing only the CC-upstream region (Fig. 1A). EMSAs indicated that the shortest fragment containing the CENP-C-like motif, KNL2^518-560^ bound to the CENP-A nucleosome but not to the H3 nucleosome (Fig. 1B). Specific binding of KNL2^518-560^ to the CENP-A nucleosome was also observed in the BLI assay (Fig. 1C and Fig. EV1). The dissociation constant (KD) of the interaction between the CENP-A nucleosome and the KNL2^518-560^ fragment was estimated to be 5.5±1.1 μM based on the BLI sensograms (Fig. EV1A). Other KNL2 fragments containing the CENP-C-like motif, KNL2^347-560^ and KNL2^456-560^ also bound to the CENP-A nucleosome (Fig. 1D and E).

Intriguingly, in the EMSAs, these longer KNL2 fragments were observed to form a complex at a lower molar ratio to the CENP-A nucleosome than that of KNL2^518-560^, suggesting more stable CENP-A nucleosome binding (Fig. 1B, D, and E). In our BLI assays, KNL2^347-560^ and KNL2^456-560^ showed stronger interactions with the CENP-A nucleosome than that of KNL2^518-560^. The KD values for the binding of KNL2^347-560^ and KNL2^456-560^ to the CENP-A nucleosome were 16.4±5.9 nM and 22.0±1.8 nM, respectively (Fig. EV1B and EV1C), whereas the KD of KNL2^518-560^ was 5.5±1.1 μM (Fig. EV1A). However, KNL2^456-516^ ^(CC-upstream)^ lacking the CENP-C like motif did not exhibit specific binding to the CENP-A nucleosome (Fig. 1F and EV1D) and we have previously shown that the SANTA alone does not specifically bind to the CENP-A nucleosome (Hori *et al*., 2017). These data suggest that the N-terminal extended region from the CENP-C-like motif (aa 347-455) contributed to the stable binding of KNL2 to the CENP-A nucleosome, although this region doesn’t directly bind to the CENP-A nucleosome. The similar affinities of KNL2^347-560^ and KNL2^456-560^ for the CENP-A nucleosome implied that SANTA would not be involved in the CENP-A nucleosome–KNL2 interaction, which is consistent with the previous observation that this domain is dispensable for centromere localization of KNL2 (Fig. EV1B and EV1C) (Hori *et al*., 2017). Thus, we concluded that only the CENP-C-like motif (KNL2^518-560^) is a sufficient region for the specific interaction between ggKNL2 and the CENP-A nucleosome, and the CC-upstream region was suggested to stabilize ggKNL2-CENP-A nucleosome binding.

### Cryo-EM structure of the CENP-A nucleosome complexed with CENP-C-like motif of ggKNL2

To obtain further insight into the binding of ggKNL2 to the CENP-A nucleosome, we performed cryo-EM single-particle analysis focusing on the CENP-C-like motif. The CENP-A nucleosome in complex with ggKNL2 was reconstituted in vitro using synthetic KNL2 peptide containing amino acid residues (aa) 517-560, KNL2^517-560^ (Fig. 2A) and subjected to cryo-EM single-particle analysis (Fig. EV2). Three-dimensional (3D) reconstruction of the CENP-A-KNL2^517-560^ complex was carried out at 3.42 Å resolution (Table 1 and Fig. EV2).

**Figure 2.**
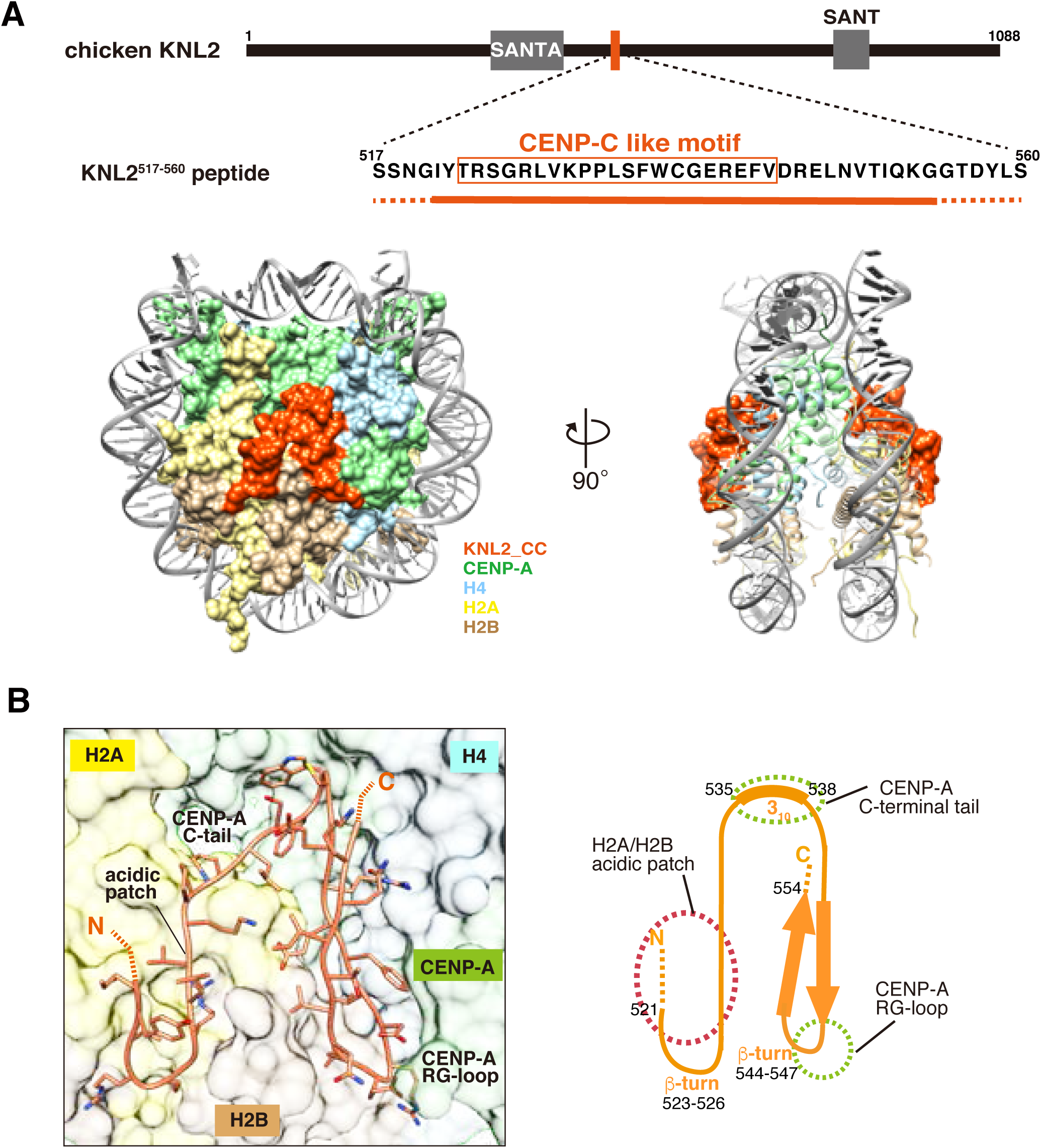
Cryo-EM structure of the CENP-C like motif of ggKNL2 bound to the CENP-A nucleosome A) Schematic diagram showing functional elements in chicken KNL2 (ggKNL2). Amino acid sequence of the chicken KNL2^517-560^ peptide (aa 517–560) used for cryo-EM analysis is presented. The region observed in Cryo-EM structure is underlined (aa 521-554). The conserved CENP-C like motif is enclosed in the orange box. Lower panels show the surface models of the CENP-A nucleosome in complex with the KNL2^517-560^ peptide. The side views of the complex along the two-fold axis are shown. DNA is presented using the cartoon model. The surface corresponding to each molecule in the complex is color-coded, as indicated in the figure. **B)** The left panel shows the stick model of the structure of the KNL2^517-560^ peptide bound to the CENP-A nucleosome. Schematic representation of the KNL2^517-560^ peptide structure is shown in the right panel. Dot circles indicate the recognition sites for the H2A/H2B acidic patch, the CENP-A C-terminal region, and the CENP-A RG-loop. See also Fig. EV2.

**Table 1.** Statistics for Cryo-EM Single Particle Image Analysis and Structure Refinement.

|  |  |
| --- | --- |
| Cryo-EM density map | KNL2_CC - CA nucleosome (3.42 Å)<br>(EMD-33666)<br>(PDB 7Y7I) |
| Data collection and processing |  |
| Sample | KNL2_CENP-C like motif peptide bound to the CENP-A nucleosome |
| Magnification | 50,000 |
| Voltage (kV) | 300 |
| Electron exposure (e<sup>-</sup>/Å<sup>2</sup>) | 40 |
| Defocus range (μm) | -1.0 to -2.5 |
| Pixel size (Å) | 0.98 |
| Symmetry | C2 |
| Initial particle images (no.) | 10,331,795 |
| Final particle images (no.) | 270,464 |
| Map resolution (Å) | 3.42 |
| FSC<sup>a</sup> threshold | 0.143 |
| Refinement |  |
| CCmap_model | 0.82 |
| Model compositions |  |
| Non-hydrogen atoms | 12447 |
| Protein residues | 824 |
| Nucleic acids | 286 |
| R.m.s. deviations<sup>b</sup> |  |
| Bond lengths (Å) | 0.004 |
| Angles (°) | 0.577 |
| Ramachandran plot |  |
| Favored (%) | 98.13 |
| Allowed (%) | 1.87 |
| Outliers (%) | 0.00 |
| Rotamer outliers (%) | 0.29 |
| Clash score | 9.14 |
| Cβ outliers (%) | 0.00 |
| CaBLAM outliers (%) | 1.28 |

a) FSC, Fourier shell correlation
b) R.m.s. deviations, root-mean-square deviations

The unambiguous cryo-EM map identified the KNL2 peptide region between residues 521 and 554 bound to each side of the symmetrical CENP-A nucleosome surfaces (Fig. 2A). The KNL2^517-560^ peptide adopts a well-defined structure on the CENP-A nucleosome and forms a stable interface with the CENP-A nucleosome (Fig. 2A and 2B). The cryo-EM structure clearly demonstrated that KNL2 shares the CENP-A nucleosome binding mode with CENP-C (Fig. EV3A-B) (Arimura *et al*, 2019; Ariyoshi *et al*., 2021; Kato *et al*, 2013b), which is consistent with the sequence similarity among their conserved CENP-C motifs (Fig. 1A). The KNL2^517-560^ peptide recognizes the H2A/H2B acidic patch using the conserved arginine residue in the N-terminal region (Fig. 2B and EV3C), while the conserved F533 and W544 of KNL2 located in a 310 helix are responsible for the recognition of the C-terminal tail of CENP-A (Fig. 2B and EV3D). As observed in the cryo-EM structure of the chicken CENP-C bound to the CENP-A nucleosome, the KNL2^517-560^ peptide makes a contact with the RG-loop of CENP-A using its C-terminal region (Fig. 2B). Furthermore, the continuous cryo-EM map at a higher resolution allowed us to identify a two-stranded antiparallel β-sheet structure in the C-terminal region of KNL2^517-560^, which was not identified in the cryo-EM structure of the chicken CENP-C bound to the CENP-A nucleosome (Fig. 2B and EV3B) (Ariyoshi *et al*., 2021). The β-turn structure (aa 544-547) in the KNL2^517-560^ peptide provided an interface with the RG-loop of CENP-A.

### CENP-C-like motif of ggKNL2 recognizes the acidic patch of histone H2A/B in the CENP-A nucleosome

As described above, the CENP-C-like motif (aa 523-543) in ggKNL2 interacts with the acidic patch of histone H2A/B (D90, E91 in H2A) in the CENP-A nucleosome (Fig. 3A), similar to the observation in the structure of the CENP-A nucleosome bound to human, rat, or chicken CENP-C (Ali-Ahmad *et al*, 2019; Allu *et al*, 2019; Ariyoshi *et al*., 2021; Kato *et al*., 2013b). Arginine 527 (R527) of ggKNL2 interacts with the D90 and E91 residues in H2A of the CENP-A nucleosome (Fig. 3A), as observed in R659 of chicken CENP-C or R522 of human CENP-C which also binds to the same site (Fig. EV3C) (Ariyoshi *et al*., 2021). To test the contribution of R527 of ggKNL2 to CENP-A nucleosome binding, we prepared a recombinant KNL2^518-560^ protein with an R527A mutation and examined its binding affinity for CENP-A nucleosome using EMSA. As shown in Fig. 3B, we observed a clear band shift for the CENP-A nucleosome complex with the wild-type KNL2^518-560^ but not for the mutant KNL2^518-560^. BLI analyses using the mutant KNL2^518-560^ and the CENP-A nucleosome showed significantly lower wavelength shifts compared to those of the wild-type protein (Fig. 3C and EV4A). Thus, the R527A mutation caused a reduction in CENP-A nucleosome binding of KNL2^518-560^.

**Figure 3.**
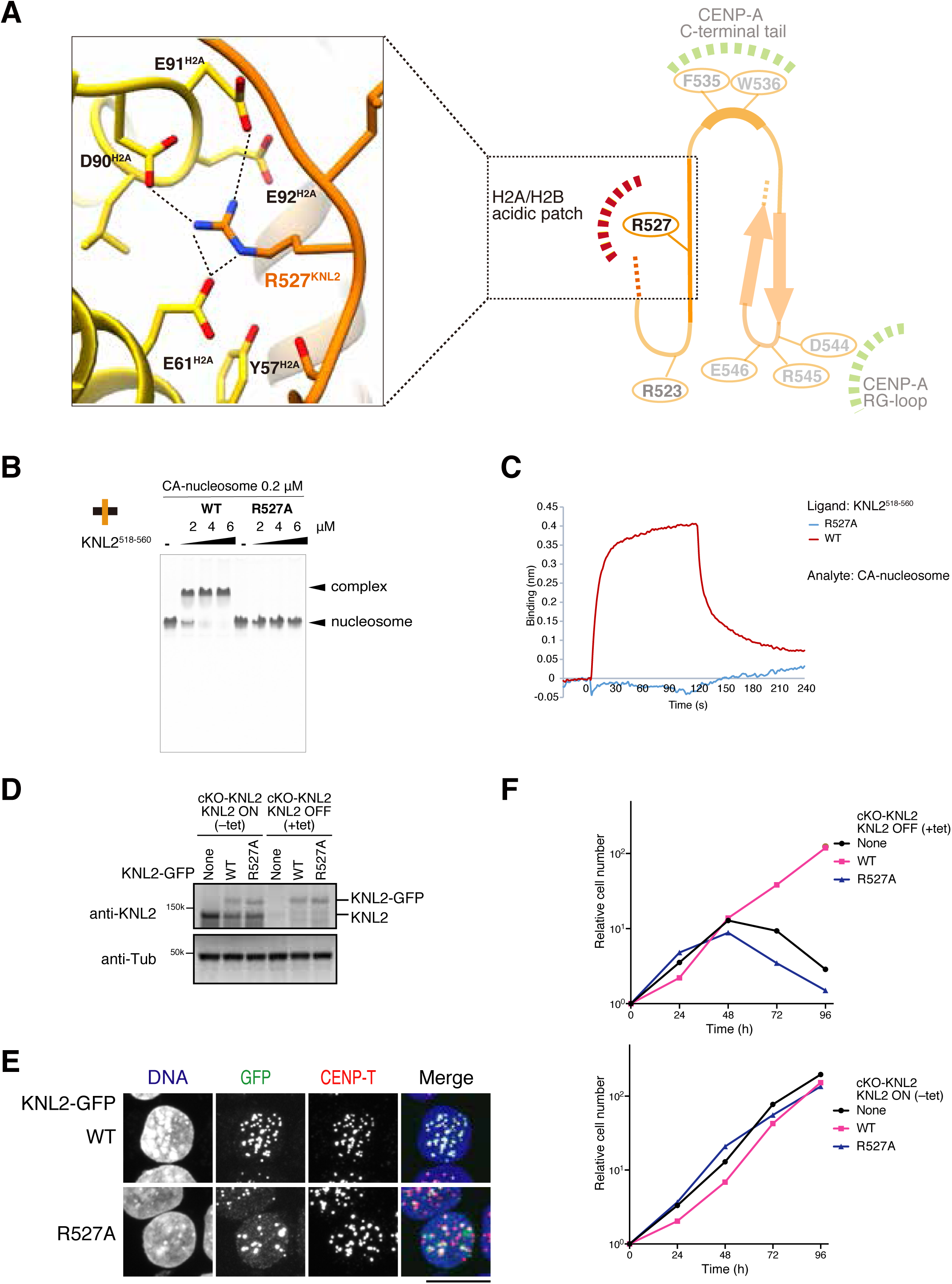
ggKNL2 recognizes the acidic patch of histone H2A/H2B A) The magnified view of the H2A/H2B binding site of KNL2. R527 of ggKNL2 forms hydrogen bonds with the H2A acidic residues. Side chains of the key residues are indicated as a stick model. Dotted lines represent the possible hydrogen bonds (<3.0 Å). **B)** EMSA to examine contribution of the ggKNL2 R527 residue to the CENP-A nucleosome binding. **C)** Biolayer interferometry analyses of the CENP-A nucleosome binding of GST-KNL2^518-560^ WT or GST-KNL2^518-560^ ^R527A^ mutant. Each GST-fused KNL2 fragment was immobilized onto anti-GST prob. 750 nM of CENP-A nucleosome was used as an analyte. **D)** Expression of the WT or R527A mutant GFP-KNL2 in KNL2 conditional knock out chicken DT40 cells (cKO-KNL2). Expression of full-length KNL2 WT was conditionally turned off by tetracycline (tet) addition in cKO-KNL2 cells (None). GFP-fused KNL2 WT or R527A mutant was stably expressed in the cKO-KNL2 cells. These cells were cultured in the presence or absence of tet (+tet: KNL2 OFF or −tet: KNL2 ON) for 48 h. α-Tubulin (Tub) was probed as a loading control. **E)** Localization analyses of GFP-fused KNL2 WT or R527A mutant (green). CENP-T was stained as a kinetochore marker (red), and DNA was stained using DAPI (blue). Scale bar indicates 10 μm. **F)** Growth of the cKO-KNL2 cells expressing GFP-fused KNL2 WT or R527A mutant. The upper panel shows examined cell numbers at the indicated time after tet addition (+tet: KNL2 OFF). The cell numbers were normalized to those at 0 h for each line and plotted as relative cell number. The lower panel shows examined untreated cells (-tet: KNL2 ON). Parental cKO-KNL2 chicken DT40 cells (None) were also examined. See also Figs. EV3A-C and EV4A.

To evaluate the importance of this residue in the KNL2 function, we introduced KNL2^R527A^-GFP into KNL2 conditional knockout chicken DT40 cells (cKO-KNL2 cells). In cKO-KNL2 cells, two alleles of the KNL2 coding region were disrupted, and the KNL2 cDNA was expressed under the control of the tetracycline (tet)-responsive promoter (Hori *et al*., 2017). Therefore, endogenous KNL2 expression could be turned off upon tet addition in this cell line. When GFP-fused ggKNL2 cDNA was expressed in the cKO-KNL2 cells after tet addition, KNL2 expression was replaced with KNL2-GFP expression. After we confirmed the replacement of endogenous KNL2 with KNL2-GFP (wild-type: WT or KNL2^R527A^ mutant) using immunoblot analysis (Fig. 3D), the localization of KNL2-GFP was examined. WT KNL2-GFP colocalized with the centromere marker CENP-T in the cKO-KNL2 interphase cells; however, KNL2^R527A^-GFP did not localize to the centromeres in the interphase cells (Fig. 3E). The KNL2^R527A^ mutant protein formed non-centromeric nuclear bodies in the interphase cells, which was caused by KNL2 lacking the entire CENP-C-like motif (Hori *et al*., 2017). Furthermore, the expression of WT KNL2-GFP suppressed growth defects in the cKO-KNL2 cells, whereas the expression of KNL2^R527A^-GFP did not suppress growth defects in the cKO-KNL2 cells, in the presence of tet (Fig. 3F). These findings indicate that the binding of ggKNL2 to CENP-A is critical for cell viability. Thus, the interactions between R527 of KNL2 and the histone residues in the acidic patch of H2A/B facilitate the complex formation of the KNL2-CENP-A nucleosome, which is essential for its centromere localization and function in DT40 cells.

### CENP-C like motif of ggKNL2 binds to the C-terminal tail of the CENP-A nucleosome

In addition to the acidic patch of H2A/B, the CENP-A C-terminal tail, which is one of the unique structural features of CENP-A, is recognized via the conserved residues F535 and W536 in the CENP-C-like motif of ggKNL2 in our cryo-EM structure (Fig. 2B and 4A). F535 and W536, which are ggKNL2 counterparts of Y667 and W668 in the CENP-C motif of ggCENP-C, or W530 and W531 in the central domain of human CENP-C (Fig. EV3D), contact CENP-A residues in the C-terminal tail. In particular, the side chain of W536 of ggKNL2 forms a stacking interaction with the side chain of R131 of CENP-A (Fig. 4A). The CENP-A C-terminal tail binding mode is well conserved between ggKNL2 and human or chicken CENP-C (Ali-Ahmad *et al*., 2019; Allu *et al*., 2019; Ariyoshi *et al*., 2021; Kato *et al*., 2013b). The side chain arrangements of these aromatic residue pairs on the CENP-A nucleosome are superposed well (Fig. EV3D).

**Figure 4.**
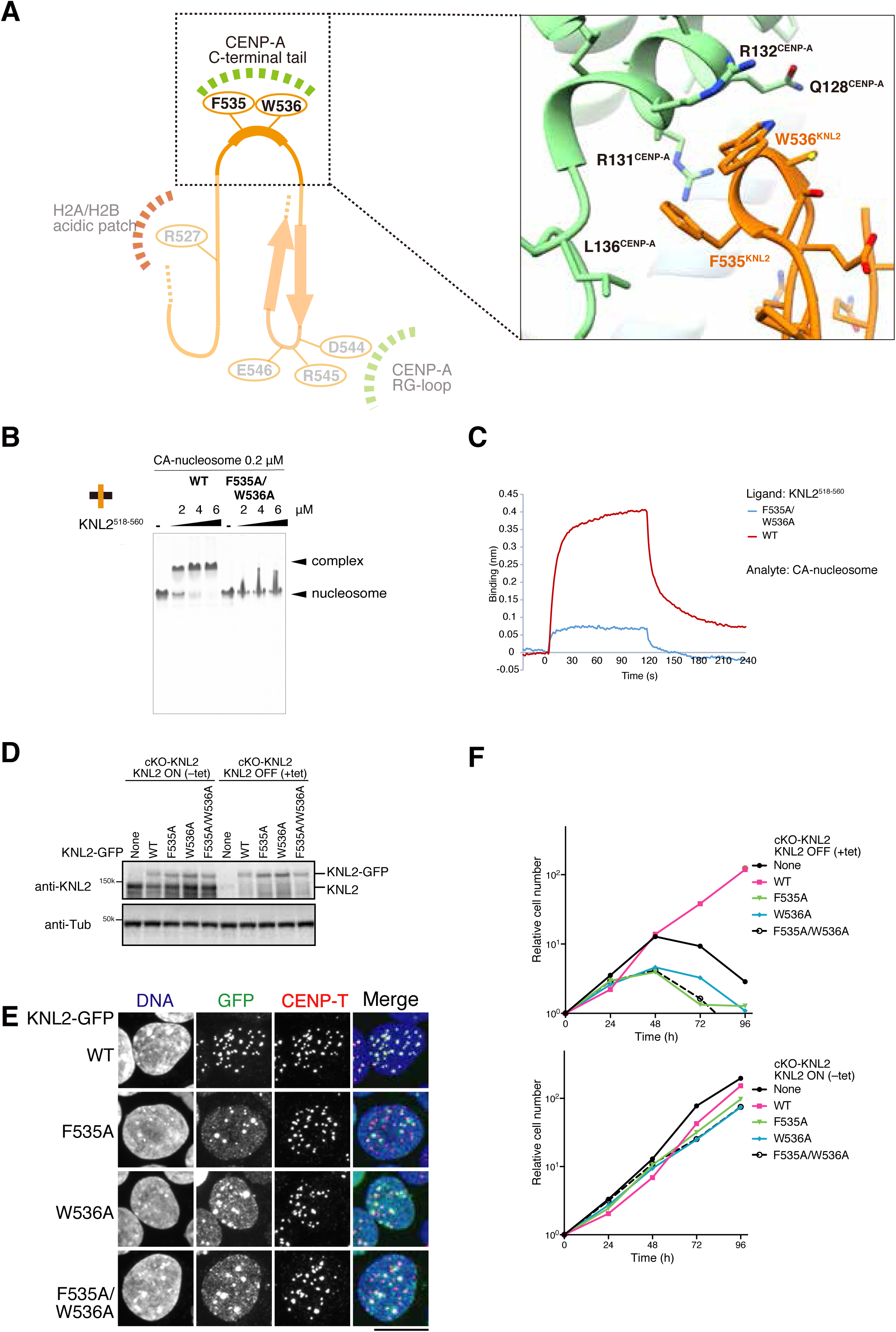
ggKNL2 recognizes the C-terminal tail of CENP-A A) Magnified view of the interface between the CENP-A C-terminal tail and the KNL2^517-560^ peptide. Side chains of the key residues are indicated as a stick model. **B)** EMSA to examine contribution of the F535 and W536 residues in ggKNL2 to the CENP-A nucleosome binding. **C)** Biolayer interferometry analyses of the CENP-A nucleosome binding of GST-KNL2^518-560^ WT or GST-KNL2^518-560^ F535/W536 mutant. Each GST-fused KNL2 fragment was immobilized onto anti-GST prob. 750 nM of CENP-A nucleosome was used as an analyte. **D)** Expression of GFP-KNL2 WT or CENP-A C-terminal tail binding mutants in cKO-KNL2 cells. Expression of full-length KNL2 WT was conditionally turned off by tetracycline (tet) addition to the cKO-KNL2 cells (None). GFP-fused KNL2 WT, F535A, W536A or F535A/W536A was stably expressed in the cKO-KNL2 cells. These cells were cultured in the presence or absence of tet (+tet: KNL2 OFF or −tet: KNL2 ON) for 48 h. α-Tubulin (Tub) was probed as a loading control. **E)** Localization analyses of GFP-fused KNL2 WT and F535A, W536A or F535A/W536A mutant (green). CENP-T was stained as a kinetochore marker (red), and DNA was stained using DAPI (blue). Scale bar indicates 10 μm. **F)** Growth of the cKO-KNL2 cells expressing GFP-fused KNL2 WT or each CENP-A C-terminal tail binding mutant. The upper panel shows examined cell numbers at the indicated time after tet addition (+tet: KNL2 OFF). The cell numbers were normalized to those at 0 h for each line and plotted as relative cell number. The lower panel shows examined untreated cells (-tet: KNL2 ON). Parental cKO-KNL2 chicken DT40 cells (None) were also examined. See also Figs. EV3A-B, EV3D and EV4B.

To evaluate the contribution of the F535 and W536 residues of ggKNL2 to CENP-A nucleosome binding, we performed CENP-A nucleosome binding assays using recombinant KNL2^518-560^ with F535A_W536A double mutations. The results of the EMSA assays indicated that this mutant did not bind to the CENP-A nucleosome (Fig. 4B). The reduced affinity for the CENP-A nucleosome caused by the F535A_W536A mutation was also confirmed using BLI analysis (Fig. 4C and EV4B). These biochemical data indicate that the F535 and W536 residues are essential for CENP-A nucleosome binding by ggKNL2.

We also examined the significance of F535 and W536 of ggKNL2 in chicken DT40 cells. GFP-fused KNL2^F535A^, KNL2^W536A^, and KNL2 ^F535A_W536A^ were introduced into the cKO-KNL2 cells (Fig. 4D). As observed in cells expressing KNL2^R527A^, each mutant protein formed non-centromeric nuclear bodies and did not properly localize to the centromeres in the interphase DT40 cells (Fig. 4E). Furthermore, unlike WT KNL2, expression of each of these mutant KNL2 proteins in cKO-KNL2 cells did not suppress the growth defects in the cKO- KNL2 cells in the presence of tet (Fig. 4F). Considering these results, the acidic patch of H2A/B and the C-terminal tail of CENP-A are critical elements for CENP-A nucleosome recognition, not only by the central domain or CENP-C-like motif in CNEP-C, but also by the CENP-C-like motif of ggKNL2.

### ggKNL2 associates with the RG-loop of the CENP-A nucleosome to distinguish between H3 and CENP-A nucleosome

In addition to the C-terminal tail, the RG-loop between helices α1 and α2 is a unique structural feature of CENP-A in comparison to that of the canonical histone H3 (Fang *et al*, 2015; Tachiwana *et al*., 2011). Our cryo-EM structure of the CENP-A nucleosome- KNL2^517-560^ complex revealed the interaction between the CENP-A RG-loop and ggKNL2. In the complex, the C-terminal part of KNL2^517-560^ (aa 539-553) forms a β-sheet, and the β-turn region (aa 544-547) was found to be located in the vicinity of the RG-loop in CENP-A (Fig. 2B). The side chain of D544 in the β-turn is supposed to make a contact with the side chain of R80 in the RG-loop of CENP-A (Fig. 5A). Indeed, KNL2^518-560^ with the D544A mutation did not exhibit CENP-A nucleosome binding in either the EMSA or BLI assay (Fig. 5B and 5C). Furthermore, we examined the functional significance of the D544 residue of KNL2 in DT40 cells. GFP-fused KNL2^D544A^ was introduced into the cKO-KNL2 cells (Fig. 5D). As observed in cells expressing other CENP-A binding site mutants, ggKNL2, KNL2^D544A^ formed non-centromeric nuclear bodies and did not properly localize to centromeres in the interphase cells (Fig. 5E). In addition, the expression of the D544A mutant in cKO-KNL2 cells did not suppress growth defects in the cKO-KNL2 cells in the presence of tet (Fig. 5F).

**Figure 5.**
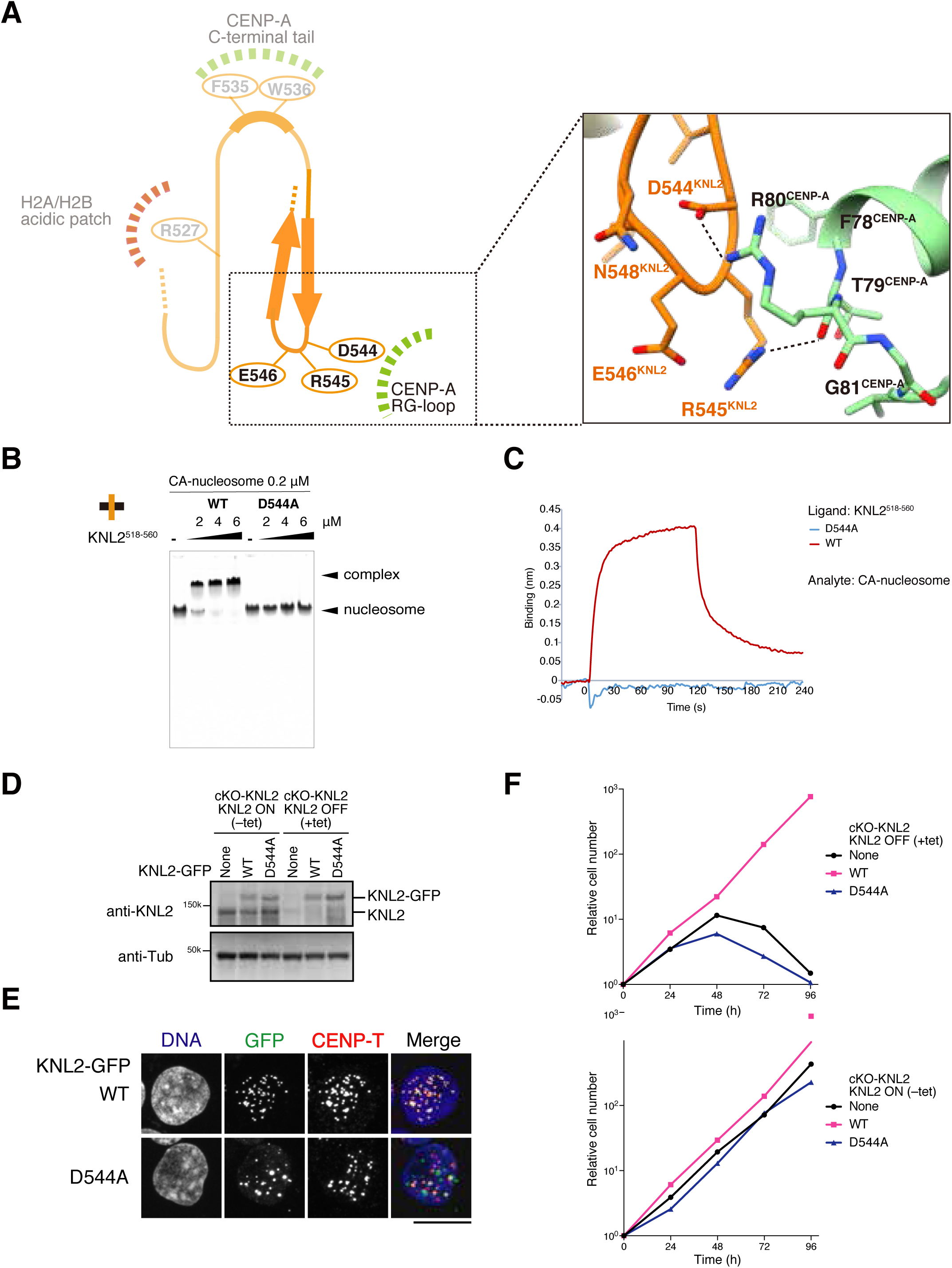
The D544 residue of ggKNL2 associates with the RG-loop of the CENP-A nucleosome A) Magnified view of the interface between the CENP-A RG-loop and ggKNL2. The β-turn structure of KNL2 provided the interface with the CENP-A RG-loop. Side chains of the key residues are indicated as a stick model. **B)** EMSA to examine the binding affinities of WT KNL2^518-560^ or KNL2^518-560^ D544A mutant to the CENP-A nucleosome. **C)** Biolayer interferometry analyses of the CENP-A nucleosome binding of GST-KNL2^518-560^ WT or GST-KNL2^518-560^ D544A mutant. Each GST-fused KNL2 fragment was immobilized onto anti-GST prob. 750 nM of CENP-A nucleosome was used as an analyte. **D)** The expression of WT ggKNL2 was conditionally turned off by tet addition in cKO-KNL2 cells (None). GFP- fused WT ggKNL2 and its D544A mutant were stably expressed in cKO-ggKNL2 cells. These cells were cultured in the presence or absence of tet (+tet: KNL2 OFF or -tet: KNL2 ON) for 48 h. α-Tubulin (Tub) was probed as a loading control. **E)** Localization of GFP- fused KNL2 WT and D544A mutant (green). CENP-T was stained as a kinetochore marker (red), and DNA was stained using DAPI (blue). Scale bar indicates 10 μm. GFP and CENP-T signals on kinetochores in mitotic cells were quantified in each cell line. **F)** Growth of the cKO-KNL2 cells expressing GFP-fused KNL2 WT and D544A mutant. The upper panel shows examined cell numbers at the indicated time after tet addition (+tet: KNL2 OFF) and were normalized to those at 0 h for each line. The lower panel shows examined untreated cells (-tet: KNL2 ON). Parental cKO-KNL2 cells (None) were also examined. See also Fig EV4C.

Considering these results, we conclude that D544 of ggKNL2 contributes to CENP-A nucleosome recognition by interacting with the CENP-A RG-loop. This finding is consistent with previous observations suggesting that the CENP-A RG-loop binds ggCENP-C (Ariyoshi *et al*., 2021). Notably, downstream regions of the conserved canonical CENP-C/CENP-C-like motifs play a key role in CENP-A recognition of both ggKNL2 and ggCENP-C.

### CENP-A localization was reduced in the cells without the KNL2-CENP-A interaction in chicken DT40 cells

If KNL2 does not localize to centromeres, a functional Mis18 complex cannot be formed on centromeres and new CENP-A incorporation does not occur (Hori *et al*., 2017). Therefore, we examined CENP-A levels at the kinetochores in the cKO-KNL2 DT40 cells expressing GFP-fused WT or each CNEP-A binding site mutant of ggKNl2: KNL2-GFP, KNL2^R527A^-GFP, KNL2^F535A^-GFP, KNL2^W536A^-GFP, KNL2^F535A_W536A^-GFP, or KNL2^D544A^-GFP (Fig. 6A and B). As shown in Fig. 6, CENP-A levels at the kinetochores in the cKO-KNL2 cells expressing mutant KNL2 (for all mutants) were significantly reduced compared to those in the cKO- KNL2 cells expressing WT KNL2. These findings indicate that abrogated binding of KNL2 to the CENP-A nucleosome reduces CENP-A incorporation through dysfunction of the Mis18 complex in chicken DT40 cells.

**Figure 6.**
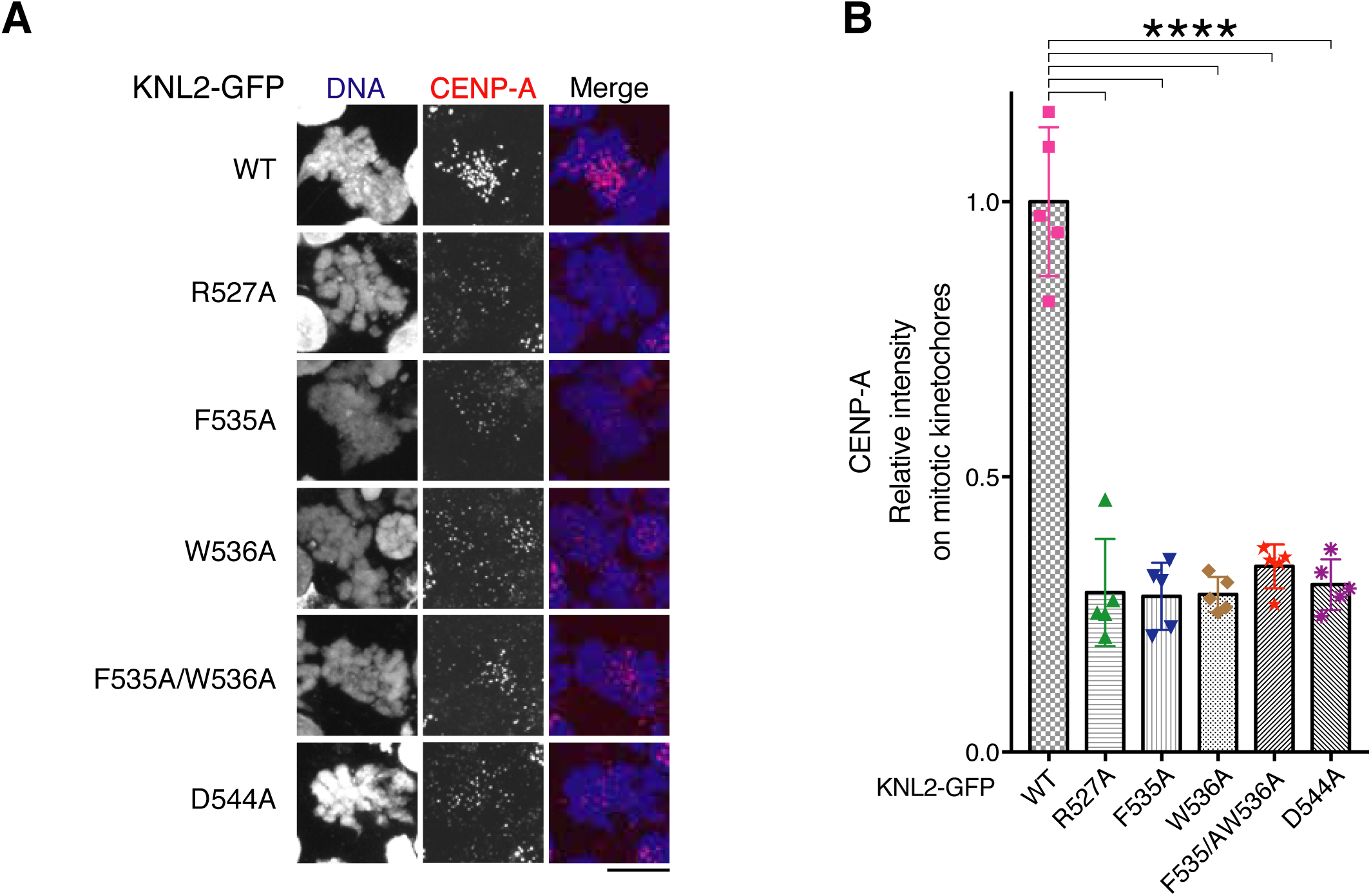
Binding of ggKNL2 to the CENP-A nucleosome via the CENP-C like motif is required for new CENP-A deposition. A) Immunostaining with anti-CENP-A antibody (red) in cKO-KNL2 cells (+tet: KNL2 OFF) expressing GFP-fused KNL2 WT or each CENP-A nucleosome binding mutant. Representative images of mitotic cells were presented. DNA was stained with DAPI (blue). Scale bar indicates 10 µm. **B)** The levels of CENP-A in mitotic cells of each mutant line were evaluated by normalizing to those in WT. Bar graph indicates mean with SD (n = 5; ****, P < 0.0001, unpaired t-test, two-tailed).

### Mitotic kinetochore localization of ggKNL2 does not depend on the CENP-A binding, but on CENP-C in chicken DT40 cells

ggKNL2 localizes to centromeres throughout the cell cycle in chicken DT40 cells (Hori *et al*., 2017). Our structural data indicated that the CNEP-C-like motif of ggKNL2 and the CNEP-C motif of ggCENP-C bind to the same surface of the CENP-A nucleosome. Such a structural similarity between ggKNL2 and ggCENP-C raises the question of whether ggKNL2 localizes to centromeres via CENP-A nucleosome interaction in the presence of ggCENP-C. Previously, we demonstrated that the centromere localization of ggCENP-C via CENP-A nucleosome binding was facilitated by CDK1 dependent phosphorylation and occurred only in mitotic cells but not in interphase cells (Ariyoshi *et al*., 2021; Watanabe *et al*., 2019). These previous findings imply that the localization dependency of ggKNL2 might differ between interphase and mitotic cells. Although we showed that centromere localization of ggKNL2 depended on the binding of the CENP-A nucleosome in chicken interphase cells (Fig. 3-5), this might not be the case in mitotic cells. To test this hypothesis, we examined KNL2 localization in the cKO-KNL2 DT40 cells expressing WT KNL2-GFP or KNL2^R527A^-GFP. As shown in Fig. 7A, KNL2 ^R527A^-GFP did not colocalize with interphase centromeres, which is consistent with the data shown in Fig. 3E. However, KNL2^R527A^-GFP perfectly colocalized to the mitotic centromeres in these cells (Fig. 7A), indicating that ggKNL2 localization to the mitotic centromeres does not involve binding to the CENP-A nucleosome.

**Figure 7.**
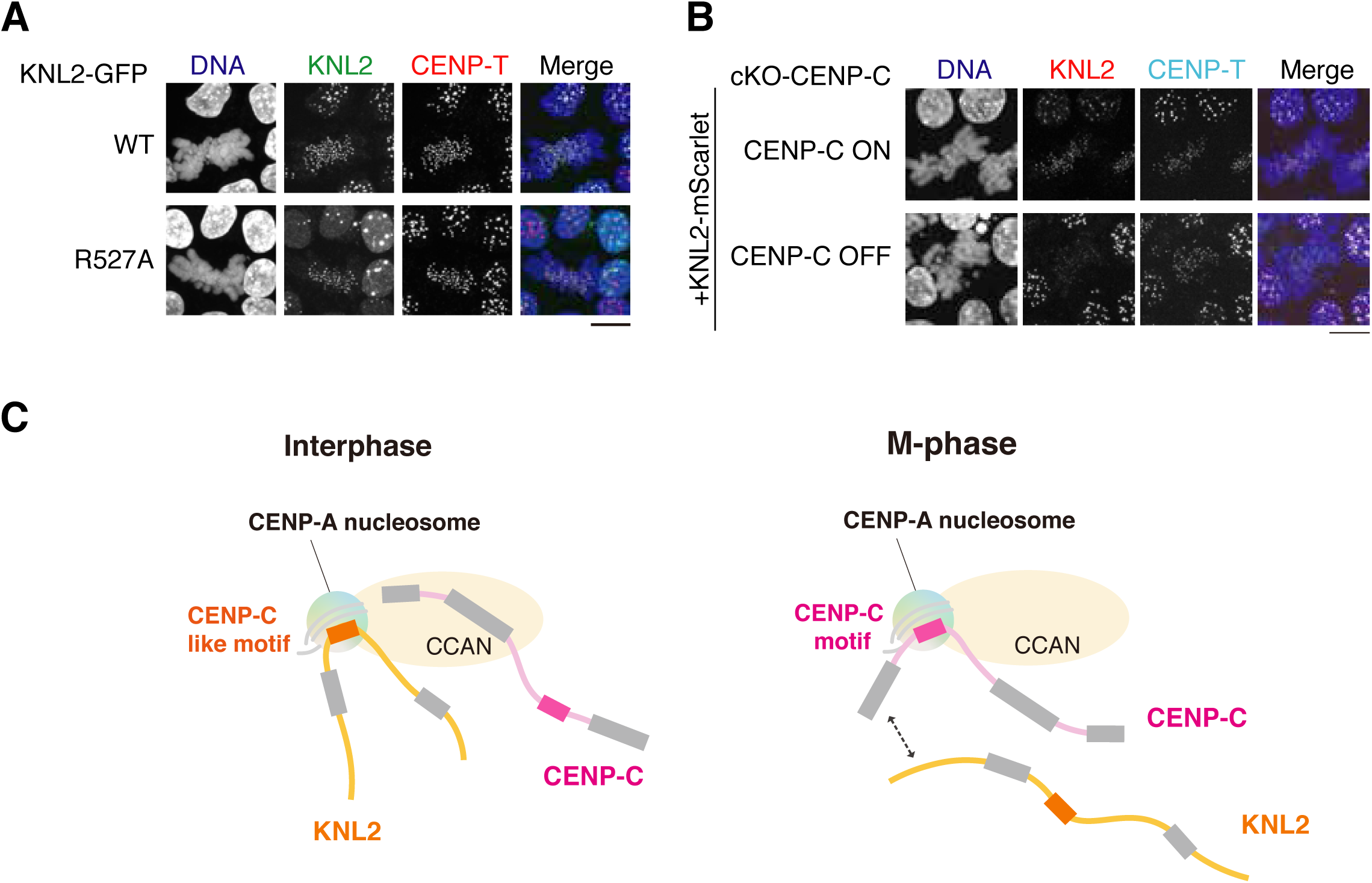
Centromere localization mode of ggKNL2 is altered during cell cycle progression A) Localization analysis of GFP-fused KNL2 WT or R527A mutant in cKO-KNL2 cells at 48 h after addition of tet (green). CENP-T was stained as a kinetochore marker (red). DNA was stained by DAPI (blue). Scale bar indicates 10 µm. **B)** KNL2 (red) localization in CENP-C conditional knockout chicken DT40 cells (cKO-CENP-C) with or without CENP-C expression. DNA was stained with DAPI (blue). CENP-T was stained as a kinetochore marker (light blue). Scale bar indicates 10 μm. **C)** A model of ggKNL2 localization dependency into centromeres in interphase and mitotic cells. ggKNL2 localizes at centromeres depending on the CENP-A nucleosome binding in interphase cells. In mitotic cells, centromere localization of ggKNL2 is mediated via CENP-C, not via CENP-A. CENP-C localizes at mitotic centromeres via the CNEP-A nucleosome interaction.

The next important question is how ggKNL2 localizes to the mitotic centromeres. Although ggKNL2 localization in interphase centromeres does not depend on CENP-C, KNL2 in other species is known to bind to CENP-C (Dambacher *et al*., 2012; Moree *et al*., 2011; Stellfox *et al*, 2016). Therefore, it is possible that the mitotic localization of ggKNL2 is mediated by CENP-C. To examine this possibility, we introduced the mScarlet sequence into the C-terminal end of the endogenous ggKNL2 gene alleles in the cKO-CENP-C cells using CRISPR-based genome editing (Watanabe *et al*., 2019) and then analyzed ggKNL2 localization in cells without CENP-C expression. Although interphase mScarlet-ggKNL2 signals remained almost unchanged, which is consistent with our previous report (Perpelescu *et al*., 2015), mitotic signals of mScarlet-ggKNL2 were not detected in the CENP-C knockout cells (Fig. 7B).

Considering the results of these cell biological analyses combined with the structural and biochemical data, ggKNL2 specifically binds to the CENP-A nucleosome via the CENP-C-like motif in the interphase nuclei, and this binding is critical for the KNL2 functions. However, mitotic centromere localization of ggKNl2 depended on CENP-C but not on the CENP-A nucleosome (Fig. 7C).

## Discussion

Unlike human or mouse KNL2, chicken KNL2 (ggKNL2) contains a CENP-C-like motif (Hori *et al*., 2017) (Fig. 1A). Since this motif is conserved in many species, including plants, we propose that a common ancestor of KNL2 has the motif, and it could have been lost during evolution to the mouse and human lineage. Since the CENP-C-like motif was demonstrated to be essential for the functioning of KNL2 in chicken DT40 cells (Hori *et al*., 2017), it is important to understand how the motif recognizes the CENP-A nucleosome. Therefore, in this study, we determined the structure of the CENP-A nucleosome in complex with ggKNL2. Our structural data revealed that ggKNL2 shares CENP-A nucleosome binding mode with CENP-C (Ariyoshi *et al*., 2021).

Our previous study (Ariyoshi *et al*., 2021) indicated that ggCENP-C recognizes the acidic patch of H2A/B and the C-terminal tail of CENP-A, which has also been reported in rat and human CENP-C studies (Ali-Ahmad *et al*., 2019; Allu *et al*., 2019; Kato *et al*., 2013a).

The structural features of these recognition sites were also well conserved in the KNL2- CENP-A nucleosome complex (Fig. EV3). In addition to these binding sites, we previously found that ggCENP-C binds to the RG-loop in CENP-A, which is absent in canonical H3 (Ariyoshi *et al*., 2021). However, the RG-loop interaction site of CENP-C remains unclear because of the low-resolution cryo-EM structural data of the CENP-C-CENP-A nucleosome complex. The cryo-EM structure of the KNL2-CENP-A nucleosome reported here revealed the interaction between KNL2 and the RG-loop in CENP-A and allowed us to identify the key residue of ggKNL2, which was D544 (Fig. 2 and 5). This new finding strongly suggests that the RG-loop interaction with CENP-C/CENP-C-like motifs contributes to distinguishing the CENP-A nucleosomes from the canonical histone H3 nucleosomes. β-sheet formation involving the downstream region of the CENP-C-like motif appears to be a key structural feature that provides the RG-loop recognition site of ggKNL2 essential for the CENP-A recognition. Thus, the stable CNEP-A nucleosome interaction with ggKNL2 or ggCENP-C requires an extra region, including the downstream region of the conserved CENP-C/CNEP-C-like motif. It is noteworthy that such an RG-loop-binding region has not been identified in the central domains of CENP-C, which is another CENP-A binding motif identified in human or mouse CENP-C. Furthermore, specific mutations introduced in each KNL2 key residue on the interaction interface with the CENP-A nucleosome reduced the binding of ggKNL2 to the CENP-A nucleosome and did not suppress the knockout phenotype of ggKNL2, indicating that each binding site of KNL2 to the CENP-A nucleosome is essential for stable KNL2- CENP-A interaction and KNL2 function.

We have previously demonstrated that CENP-C–CENP-A nucleosome interaction does not occur in interphase chicken DT40 cells, but CENP-C binds to the CENP-A nucleosome upon CDK1-mediated CENP-C phosphorylation during mitosis (Ariyoshi *et al*., 2021; Watanabe *et al*., 2019; Watanabe *et al*, 2022). We have also demonstrated that CDK1 mediated CENP-C phosphorylation facilitates the CENP-C–CENP-A interaction in human mitotic cells (Watanabe *et al*., 2019). However, since human CENP-C has two CENP-A binding sites (Klare *et al*, 2015) and chicken has only one site, another human CENP-A binding domain (central domain) of CENP-C may still bind to the CENP-A nucleosome in interphase cells, but chicken CENP-C may not associate with the CENP-A nucleosome in interphase cells. Recent structural studies on human constitutive centromere associated network (CCAN) and the CENP-A nucleosome suggest that CENP-N does not directly bind to the CENP-A nucleosome in the context of CCAN (Pesenti *et al*, 2022; Yatskevich *et al*, 2022), although previous studies suggested CENP-N-CENP-A nucleosome interaction via the RG-loop without other CCAN components (Carroll *et al*, 2009; Chittori *et al*, 2018; Pentakota *et al*, 2017; Tian *et al*, 2018). These recent cryo-EM data suggest that neither CENP-C nor CENP-N bind to the CENP-A nucleosome in interphase chicken DT40 cells. Thus, we propose that ggKNL2 associates with CENP-A nucleosome instead of CENP-C or CENP-N in interphase chicken DT40 cells (Fig. 7C). As human KNL2 was not shown to bind to the CENP-A nucleosome, the central domain of CENP-C, which is an additional CENP-A binding site, may be associated with the CENP-A nucleosome in human interphase cells; ggKNL2 may have functions similar to that of the human central domain in interphase cells.

Since the human central domain of CENP-C is involved in CENP-A stability via CENP-A binding (Falk *et al*, 2015), ggKNL2 might also be involved in the stabilization of the CENP-A nucleosome, in addition to CENP-A deposition via Mis18 complex formation.

During mitosis, CENP-C binds to the CENP-A nucleosome. Interestingly, the localization of ggKNL2 to the mitotic kinetochore does not depend on CENP-A, suggesting that ggKNL2 might dissociate from the CENP-A nucleosome during mitosis (Fig. 7C). We have previously demonstrated that CENP-C preferentially associates with the CENP-A nucleosome via mitosis-specific CENP-C phosphorylation. In addition to CENP-C phosphorylation, KNL2 dissociation from the CENP-A nucleosome might facilitate the CENP-C–CENP-A nucleosome interaction. After dissociation from the CENP-A nucleosome, ggKNL2 remains at the centromeres. As we observed that KNL2 localization at the mitotic centromeres depends on CENP-C, we proposed that KNL2 might associate with CENP-C during mitosis (Fig. 7C). The regulatory mechanisms by which KNL2 changes its binding partner between interphase and mitosis are still unclear. In addition, biological significance of mitotic localization of ggKNL2 has to be investigated. Since Mis18α does not localize at the mitotic centromeres (Hori *et al*., 2017), ggKNL2 may have a function different from Mis18 complex formation at mitotic centromeres. Understanding the mechanisms of KNL2 localization to mitotic centromeres and their biological significance will be an important subject in the future.

KNL2 is a 1088 aa long protein that contains several domains in addition to the CENP-C-like motif. The extreme N-terminal region (aa 1-140) binds to Mis18α/μ (Pan *et al*., 2017; Pan *et al*., 2019; Spiller *et al*, 2017), and this region is essential for the KNL2 function. Although the SANT and SANTA domains are dispensable for DT40 cell viability (Hori *et al*., 2017), the cKO-KNL2 cells expressing the KNL2 without either SANT or SANTA domains showed slower growth than that of the cKO-KNL2 cells expressing the WT KNL2, suggesting that these domains may facilitate the KNL2 function to some extent. Although the upstream region of the CENP-C-like motif, including the SANTA domain, did not directly bind to the CENP-A nucleosome, it facilitated stable KNL2-CENP-A nucleosome interaction (Fig. 1). Structure prediction of this region using Alphafold2 did not reveal a clear structure for this region except for the SNATA domain. However, it will be important to investigate how the upstream regions facilitate CENP-C-like motif-CENP-A nucleosome interaction.

In this study, using the ggKNL2 peptide containing the CENP-C-like motif and the reconstituted CENP-A nucleosome, we established the binding mode of ggKNL2 to the CENP-A nucleosome. We revealed that the stable complex structure is similar to that of the CENP-C–CENP-A nucleosome complex, which might explain why CENP-C does not bind to the CENP-A nucleosome in chicken DT40 interphase cells. We believe that our findings provide important clues for understanding the structure of CENP-A chromatin in species having the KNL2 with the CENP-C-like motif.

## Materials and Methods

### Recombinant protein expression and purification

cDNA fragments encoding the C-terminal fragments of chicken KNL2 (aa 347- 560, KNL2^347-560^; aa 347-455, KNL2^347-455^; aa 456-560, KNL2^456-560^; aa 456-516, KNL2^456-516^; aa 518-560, KNL2^518-560^) were each inserted into a pGEX-6p bacterial expression vector (Cytiva; pGEX-KNL2). The FLAG tag was fused to the C-terminus of each fragment. Each KNL2 fragment was expressed in *E. coli* Rosetta^TM^2 (DE3) (Novagen) as a glutathione S-transferase (GST)-fused recombinant protein. *E. coli* cells were grown in LB-Lennox media at 37°C until OD600 reached 0.6; protein expression was induced by addition of isopropyl-β-D-thiogalactoside (IPTG) to a final concentration of 0.2 mM, and culture was continued at 17°C overnight for the GST-KNL2^347-560^ and GST-KNL2^347-455^ fragments, and at 37°C for 2 hours for GST-KNL2^456-560^, GST-KNL2^518-560^. Each of KNL2 fragments was purified as follows.

Cells were harvested by centrifugation. The cell pellet was resuspended in GST-binding buffer (20 mM HEPES pH 7.5, 300 mM NaCl, 5% glycerol and 2 mM Dithiothreitol (DTT)) supplemented with protease inhibitor cocktail (Roche) and lysed by sonication on ice. The lysate was clarified by centrifugation and applied to glutathione Sepharose FF (Cytiva) pre-equilibrated with GST-binding buffer. After extensive column washing with GST-binding buffer, the immobilized GST-fusion protein was eluted from the column with GST-elution buffer (50 mM HEPES pH 7.5, 300 mM NaCl, 5% glycerol and 10 mM reduced glutathione) and applied to a cation or anion exchange column: a HiTrap SP HP cation exchange column (Cytiva) for GST-KNL2^347-560^ and GST-KNL2^456-560^; a Resource Q anion exchange column (Cytiva) for GST-KNL2^518-560^. Each GST-KNL2 fragment was eluted with a linear NaCl gradient from 70 mM to 700 mM, and further purified by size-exclusion column chromatography using a HiLoad Superdex 16/60 200 pg column (Cytiva) in a buffer containing 20 mM HEPES pH 7.5, 300 mM NaCl, 5% glycerol and 2 mM DTT. Peak fractions containing the GST-KNL2 fragment were combined, concentrated (typically to 3-5 mg/ml), and stored at -80°C until further use for structural and biochemical analyses. The expression vectors of KNL2 mutants were generated based on the pGEX-KNL2^518-560^ vector. The GST-KNL2 mutant proteins were expressed and purified in the similar manner to the wild type GST-KNL2^518-560^ fragment.

### Purification of histones

Chimeric chicken CENP-A (H3^1-63^-CA^α1-end^*), which N-terminal tail (aa 1-54 in chicken CENP-A) is substituted with that of canonical H3 (aa 1-63 in human H3), was used for chicken type CENP-A nucleosome reconstitution. Chimeric chicken CENP-A, H3^1-63^-CA^α1-end^* with an N-terminal His-tag was expressed in *E-coli*, produced as inclusion body and purified using Ni affinity and anion exchange column chromatography under the denatured conditions as previously described (Ariyoshi *et al*., 2021). Tag-free H3^1-63^-CA^α1-end^* was lyophilize using a vacuum centrifugal concentrator (TOMY). Histone H3.2 or H4 was prepared as previously described (Watanabe *et al*., 2019). H3^1-63^-CA^α1-end^* and His-H4 were mixed at 1:1 molar ratio in refolding buffer (10 mM Tris pH 7.5, 1 M NaCl, 1 mM EDTA and 5 mM DTT) and assembled as tetramers by three dialysis steps in which NaCl concentration was decreased stepwise to 200 mM. The tetramer was further purified by size-exclusion column chromatography using a HiLoad Superdex 16/60 200 pg columns (Cytiva) in buffer containing 10 mM Tris pH 7.5, 200 mM NaCl, 1 mM EDTA and 5 mM DTT, and concentrated using Amicon Ultra centrifugal filter (Merck). The gH3.2/H4 tetramer was prepared in the same way. Histones H2A and H2B were expressed in *E. coli* Rosseta2(DE3) as a homodimer from a pETDuet-His-SUMO-H2A/H2B plasmid which generates N-terminally 6xHis-SUMO–tagged H2A and untagged H2B. Bacteria cells were resuspended in high-salt buffer (20 mM HEPES pH 7.5, 2 M NaCl, 5% glycerol, and 5 mM TCEP) and disrupted by sonication. The clarified lysate was applied to a Ni-NTA agarose column (Qiagen). The H2A/H2B dimer was eluted from the column with high-salt buffer containing 300 mM imidazole and was incubated with SENP7 protease to remove the His-SUMO-tag.

The complex was further purified using HiLoad Superdex 16/60 75 pg and HiLoad Superdex 16/60 200 pg columns (Cytiva) in buffer containing 10 mM Tris pH 7.5, 200 mM NaCl, 1 mM EDTA and 5 mM DTT. The purified H2A/H2B dimer were concentrated using Amicon Ultra centrifugal filter (Merck).

### Reconstitution of chicken CENP-A nucleosomes

145 bp 601 DNA was expressed as previously described (Arimura *et al*, 2012; Tanaka *et al*, 2004). H2A/H2B dimers, chicken (CENP-A/H4)2 hetero-tetramers, and 601 DNA were mixed with a molar ratio of 2:1:0.8 at 0.75 mg/ml of final DNA concentration in the presence of 2 M KCl. A gradient dialysis to low salt buffer (2 - 0.2 M KCl in 10 mM Tris pH 7.5, 1 mM EDTA, 5 mM DTT) was performed at 4°C over 21 hours (Dyer *et al*, 2004). Finally, the nucleosome solution was dialysis against 10 mM Tris-HCl buffer (pH7.5) containing 50 mM NaCl, 1 mM EDTA, 5 mM DTT. Assembled nucleosomes were then uniquely positioned on the DNA by a thermal shift for 1 h at 37°C. Nucleosome formation was examined using native polyacrylamide gel electrophoresis. The H3 nucleosome was reconstituted in the same way as chicken CENP-A nucleosome. The concentration of each nucleosome sample was determined using absorbance at 260 nm.

### EMSA assays

Each of GST-KNL2 fragments was mixed with CENP-A or H3 nucleosome in binding buffer (20 mM HEPES pH 7.5, 1 mM EDTA, 100 mM NaCl, 5 mM TCEP, 5% glycerol and supplemented with protease inhibitor cocktail (Roche)) for 60 minutes on ice. The mixtures were analyzed by Native-PAGE using SuperSep 5-20% gels (FUJIFILM WAKO Chemicals). Native PAGE was typically performed at constant 200 V for 100 min at 4°C in 0.5xTAE buffer. The gels were stained with GelRed^TM^ (Biotium, Inc) and visualized by UV illumination at 260 nm.

### Bio-layer interferometry (BLI) assays

The association and dissociation between each GST-KNL2 fragment and CENP-A nucleosome was measured using the BLItz biolayer interferometer (FortéBio, Fremont, CA, USA). Each GST-fused KNL2 fragment was loaded onto the anti-GST biosensors (FortéBio) as follows: Biosensor probes were hydrated in BLI-binding buffer (20 mM HEPES-NaOH (pH 7.5), 200 mM NaCl, 2 mM TCEP, 1 mM EDTA, 0.5% CHAPS) for 10 min followed by loading 0.6 μM of each GST-fused KNL2 fragment (ligand) and incubated for 2 min. The ligand-bound sensor tips were washed with the BLI-binding buffer for 2 min. The CENP-A nucleosome (analyte) dissolved in BLI-binding buffer was loaded onto the GST-fused KNL2 fragment sensor tips at each concentration indicated in the BLI figures. The association phase (2 min) was measured followed by a dissociation phase (2min) in BLI-binding buffer. All the BLI measurements were performed at room temperature. Sensorgrams were analyzed by BIAevaluation 4.1 (BIACORE) to perform steady state affinity fitting to estimate apparent equilibrium dissociation constant.

### Cryo-EM data collection

The CENPC-motif peptide derived from KNL2 (aa 517-560, KNL2^517-560^ peptide) was commercially synthesized (Biologica Co. Ltd.) and dissolved in 10 mM Tris-HCl buffer (pH 7.5). The KNL2^517-560^ peptide and CENP-A nucleosome were mixed at a 5:1 molar ratio in 250 μl ul of binding buffer (20 mM HEPES pH 7.5, 100 mM NaCl, 0.1% CHAPS and 5 mM TCEP). The mixture was concentrated using an Amicon Ultra 0.5 ml filter unit (50 kDa molecular weight cut off, Merck). A final concentration of the complex was approximately 11.2 µM. A 2.5 µl protein solution of each CENP-A nucleosome complex was applied to Quantifoil Cu R0.6/1.0 holey carbon grids and frozen in liquid ethan using a Vitrobot IV (FEI, 4 °C and 100 % humidity). Data collection of each sample was carried out on a CRYO ARM 300 (JEOL, Japan) equipped with a cold field-emission electron gun operated at 300 kV, an Ω-type energy filter with a 20 eV slit width and a K3 Summit direct electron detector camera (Gatan, USA). An automated data acquisition program, SerialEM (Mastronarde, 2005), was used to collect cryoEM image data. Movie frames were recorded using the K3 Summit camera at a calibrated magnification of × 50,000 corresponding to a pixel size of 0.98 Å with a setting defocus range from -1.0 to -2.5 μm. The data were collected with a total exposure of 3 sec fractionated into 40 frames, with a total dose of ∼40 electrons Å^−2^ in counting mode. 9,450 movies were collected.

### Image processing and 3D reconstruction

Motion correction was carried out by MotionCor2 (Zheng *et al*, 2017) to align all CENP-A nucleosome complex micrographs and the CTF parameters were estimated by Gctf (Zhang, 2016). All CENP-A complexes were automatically selected by Auto-picking using the Laplacian and Gaussian in RELION 3.1 (Zivanov *et al*, 2018), and they were extracted into a box of 192 × 192 pixels. 10,331,795 particles were selected after performing 2D classification by RELION 3.1. An initial 3D classification into three classes with C1 symmetry resulted in one class (629,490 particles) that was used as further 3D refinement, CTF refinement procedure and particles polishing. The map for a well-defined nucleosome structure and the KNL2^517-560^ peptide at 3.36 Å resolution was obtained after solvent mask post-processing.

Further focused 3D classification with a mask generated for the KNL2^517-560^ peptide resulted in one class (270, 464 particles). The map of the CENP-A nucleosome - KNL2^517-560^ peptide complex at 3.42 Å resolution was obtained after 3D refinement, particle polishing, CTF refinement and solvent mask post-processing with C2 symmetry.

### Model building

The crystal structure model of the nucleosome containing the chimeric H3/CENP-A (PDB entry, 5Z23) and the cryo-EM structure of the CENP-C fragment in complex with CENP-A nucleosome (PDB entry, 7BY0) were modified to generate an initial model. Except for the canonical CENP-C motif region (aa 654 - 675), the model of CENP-C was removed and the unconserved residues were manually replaced by alanine in program Coot (Emsley *et al*, 2010). The initial model was fit into the cryo-EM density of the CENP-A nucleosome - KNL2^517-560^ peptide complex using UCSF Chimera (Pettersen *et al*, 2004). The model of the KNL2^517-560^ peptide was manually replaced its sequence by that of chicken KNL2 and modified using Coot. The model for the further region of KNL2 was manually built. The model was subjected to real space refinement in PHENIX (Liebschner *et al*, 2019). Based on the EM map density, the complex model was iteratively modified in Coot and refined in PHENIX. The final model coordinates and maps were deposited in the protein data bank. The structural figures were produced using UCSF Chimera.

### Cell culture

Chicken DT40 cells were cultured in DMEM medium (Nacalai-tesque) supplemented with 10% fetal bovine serum (FBS; Sigma), 1% chicken serum (Gibco), 10 μM 2-mercaptoethanol, Penicillin/Streptomycin (final: 100 unit/ml and 100 μg/ml, respectively) (Thermo Fisher) at 38.5°C with 5% CO2. KNL2 conditional knockout cells (Hori *et al*., 2017) were used for examination of centromere localization GFP fused KNL2. Various KNL2 mutant DNAs were cloned into a pEGFP-C3 vector (Clontech), which were transfected into KNL2 knockout cells with Gene Pulser II electroporator (Bio-Rad). The transfected cells were selected in the medium containing 2 mg/ml G418 (Santa Cruz Biotechnology).

### Immunoblotting

Expression of each protein in culture cells was analyzed by immunoblotting. For whole-cell samples, DT40 cells were harvested, washed with PBS, and suspended in 1× Laemmli sample buffer (LSB; final 10^4^ cells/μl), followed by sonication and heating for 5 min at 96°C. Proteins were separated on SuperSep Ace, 5–20% (FUJIFILM WAKO Chemicals), and transferred to Immobilon-P (Merck) using HorizeBLOT (ATTO). Primary antibodies used in this study were rabbit anti-KNL2, and mouse anti-α-tubulin (Sigma). Secondary antibodies were HRP-conjugated antirabbit IgG (Jackson ImmunoResearch), HRP-conjugated antimouse IgG (Jackson ImmunoResearch). To increase sensitivity and specificity, Signal Enhancer Hikari (Nacalai Tesque) was used for all antibodies. The antibodies were incubated with the blotted membranes for 1 h at room-temperature or for overnight at 4°C. Proteins reacting with antibodies were detected with ECL Prime (Citiva) and visualized with ChemiDoc Touch (Bio-Rad). Acquired images were processed using Image Lab 5.2.1 (Bio-Rad) and Photoshop CC (Adobe).

### Immunofluorescence and image acquisition

DT40 cells expressing various GFP fused KNL2 were centrifuged onto slide glasses by the Cytospin3 centrifuge (Shandon), and fixed with 4% paraformaldehyde (PFA; Thermo Fisher) in PBS for 10 min at room temperature. The fixed cells were permeabilized in 0.5% NP-40 in PBS for 10 min at room temperature and incubated with rabbit anti-CENP-T (1: 1,000) (Hori *et al*., 2008) in 0.5% BSA in PBS for 1 h at 37°C as a primary antibody. The reacted cells were washed with 0.5% BSA (Equitech-Bio Inc) in PBS at 3 times and were incubated with Cy3-conjugated mouse anti-rabbit IgG (1:2,000; Jackson ImmunoResearch) in 0.5% BSA in PBS for 30 min at 37°C as a secondary antibody. DNA was stained with 1 μg/ml DAPI in PBS for 1 min and mounted with VECTASHIELD mounting medium (Vector Laboratories). Immunofluorescence images were acquired every 0.2 μm step of Z-slice (total 4∼8 μm thick) as a Z-stack image using a Zyla 4.2 sCMOS camera (ANDOR) mounted on a Nikon Eclipse Ti inverted microscope with an objective lens (Plan Apo lambda 100x/1.45 NA; Nikon) with a spinning disk confocal scanner unit (CSU-W1; YOKOGAWA) controlled by NIS-elements (Nikon). The images in figures are the maximum intensity projection of the Z-stack generated with NIS-elements and were processed by Photoshop CC (Adobe).

### Quantification and statistical analysis

Protein and DNA concentrations were measured in a Nanodrop 2000 (Life technologies). The fluorescence signal intensities of CENP-A on kinetochores were quantified using Imaris (Bitplane). The fluorescence signals on ∼50–100 kinetochores in each of seven cells were quantified and subtracted with mean of background signals in non-kinetochore regions. The mean of the kinetochore signals in each cell is shown. Data are shown as mean ± SD. The unpaired t test (two tailed) was used, with P < 0.05 considered significant.

## Data availability

The cryo-EM density map of the CENP-A nucleosome - KNL2^CC^ peptide complex has been deposited into the EMDataBank with accession codes EMD-33666. The atomic coordinates have been deposited in the Protein Data Bank with accession codes PDB: 7Y7I.

## Acknowledgments

The authors are very grateful to, K. Oshimo and R. Fukuoka for technical assistance. This work was supported by the grants of CREST of JST (JPMJCR21E6), JSPS KAKENHI Grant Number 22H00408, 22H04692, 21H05752, 17H06167, to TF, the China Scholarship Council (CSC), CSC number 202108050122 to JH, JSPS KAKENHI Grant Number to 21K06048 to MA, JSPS KAKENHI Grant number 25000013, Platform Project for Supporting Drug Discovery and Life Science Research (BINDS) from AMED under Grant Number JP19am0101117, the Cyclic Innovation for Clinical Empowerment (CiCLE) Grant Number JP17pc0101020 from AMED, and JEOL YOKOGUSHI Research Alliance Laboratories of Osaka University to KN.

## Author contributions

H.J. prepared materials and performed all biochemical experiments under supervision of M.A. M.A. H.J., and F.M. performed all cryo-EM experiments and K.N. helped cryo-EM single particle image. R.W. performed DT 40 experiments. T.F. supervised the entire project and T.F. and M.A. wrote the manuscript, discussing with all authors.

## Expanded View Figure legends

**Figure EV1.**
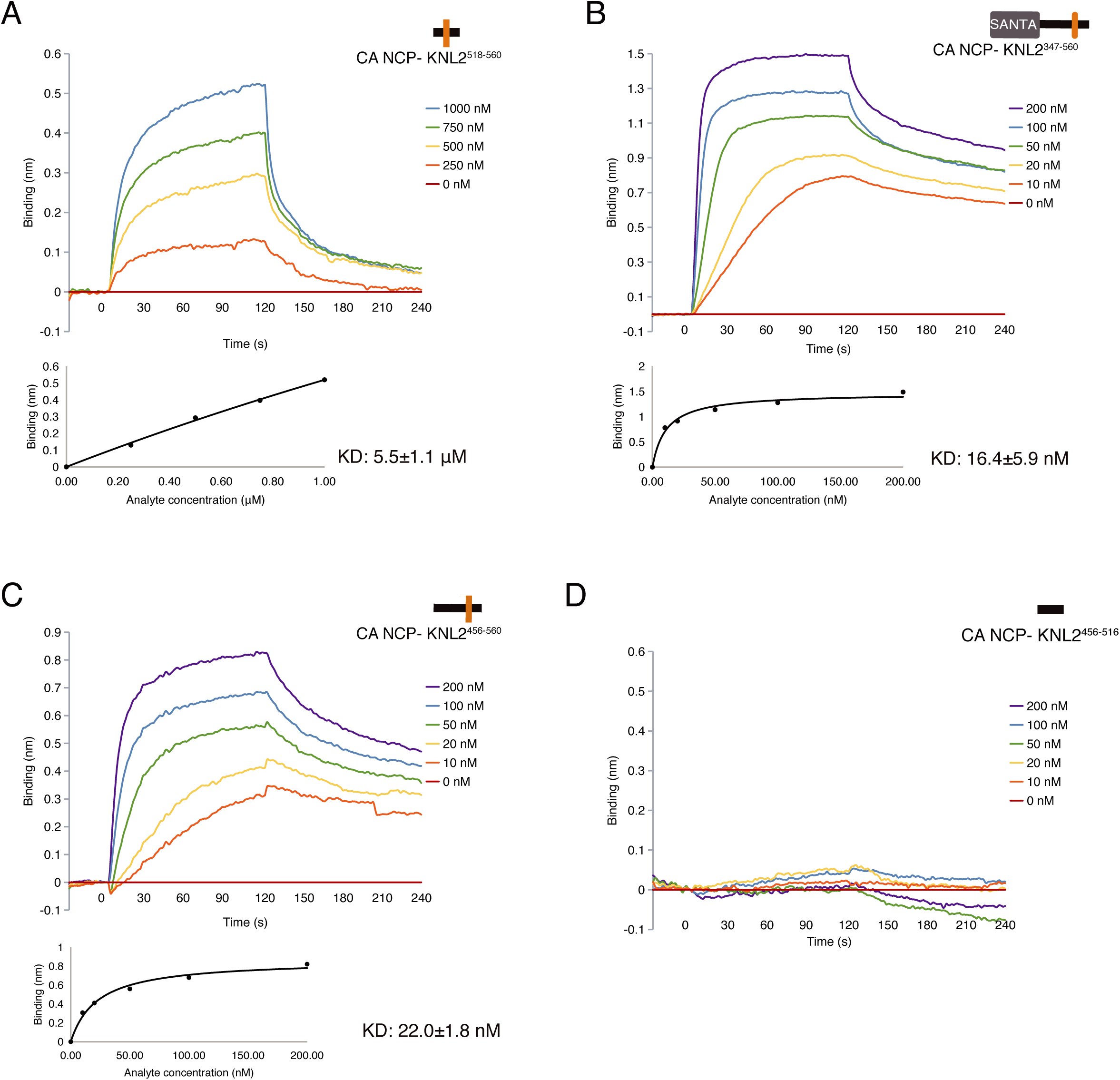
Biolayer interferometry (BLI) kinetics analyses of the interactions between CENP-A nucleosome and ggKNL2 fragments. BLI assays to examine the interaction between the GST-KNL2 fragment (immobilized ligand) and CENP-A nucleosome (analyte): **A),** GST-KNL2^518-560^ **B),** GST-KNL2^347-560^ **C),** GST-KNL2^456-560^ **D)**, GST- KNL2^456-516^. Representative BLI sensorgrams measured with analyte concentrations ranging from 0 to 1 μM in (A) or from 0 to 200 nM in (B)–(D) are plotted in the upper panel. Biolayer interferometry-derived steady-state analysis of the binding response (nm) as a function of the concentration of each KNL2 fragment is presented in the lower panel of (A) - (C). Binding assays were replicated three times. The BLI sensorgrams were measured at association time intervals of 120 s and dissociation time intervals of 120 s.

**Figure EV2.**
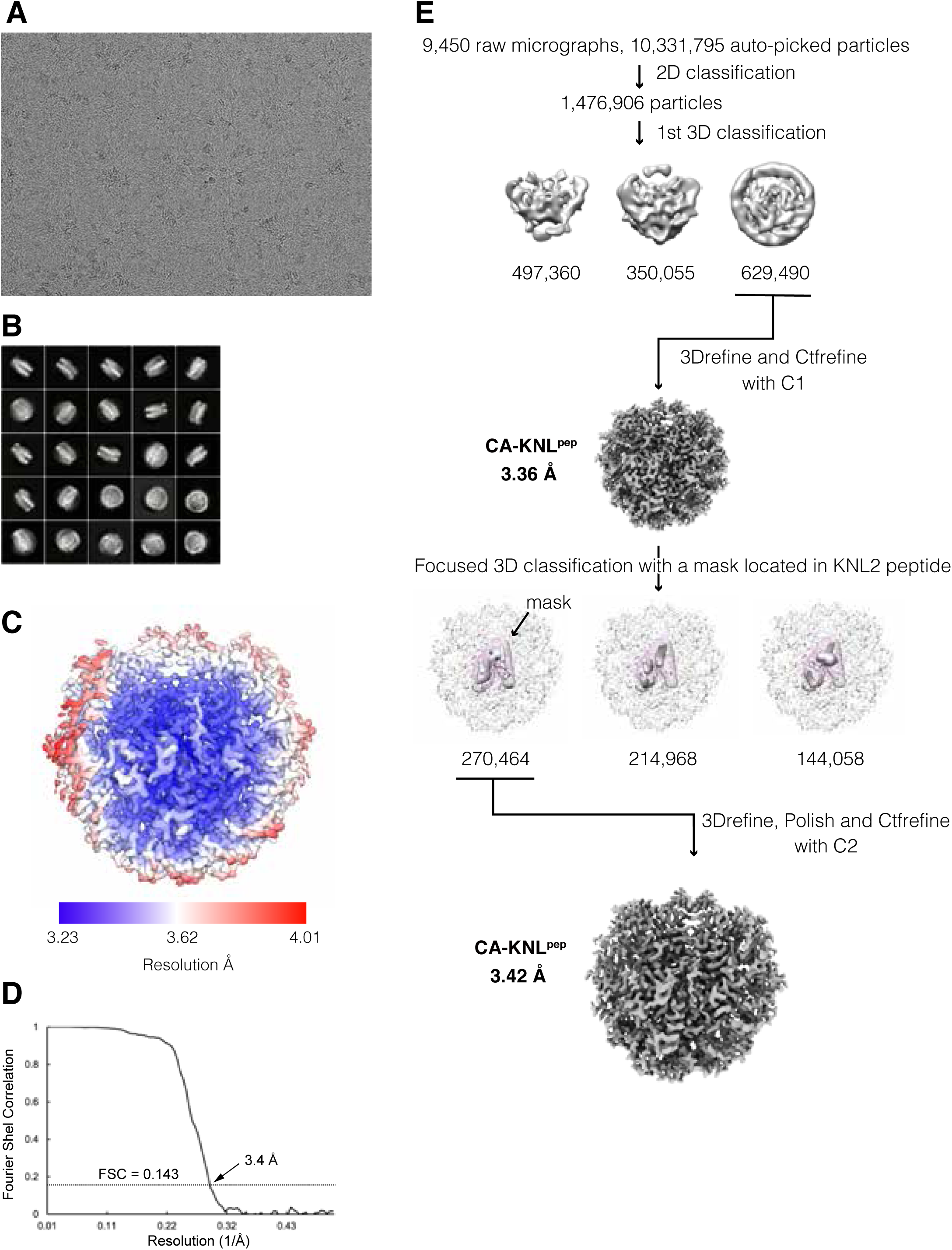
Cryo-EM single-particle image analysis of CA-ggKNL2^518-560^ complexes. A) A representative micrograph of the CENP-A nucleosome in complex with the ggKNL2^517-560^ peptide. **B)** Representative 2D class averages of the CENP-A nucleosome - KNL2^517-560^ complex. **C)** Cryo-EM density map of the chicken CA-KNL2^517-560^ complex at a 3.4 Å resolution, colored according to the local resolution estimated by RELION. **D)** Gold-standard Fourier shell correlation (FSC) curve of the Cryo-EM density map is displayed. Reported resolution (3.42 Å) was based on the FSC = 0.143 criterion. **E)** Flow chart showing the image processing pipeline for the cryo-EM single-particle image analysis of the chicken CENP-A nucelsome-KNL2^517-560^ complex.

**Figure EV3.**
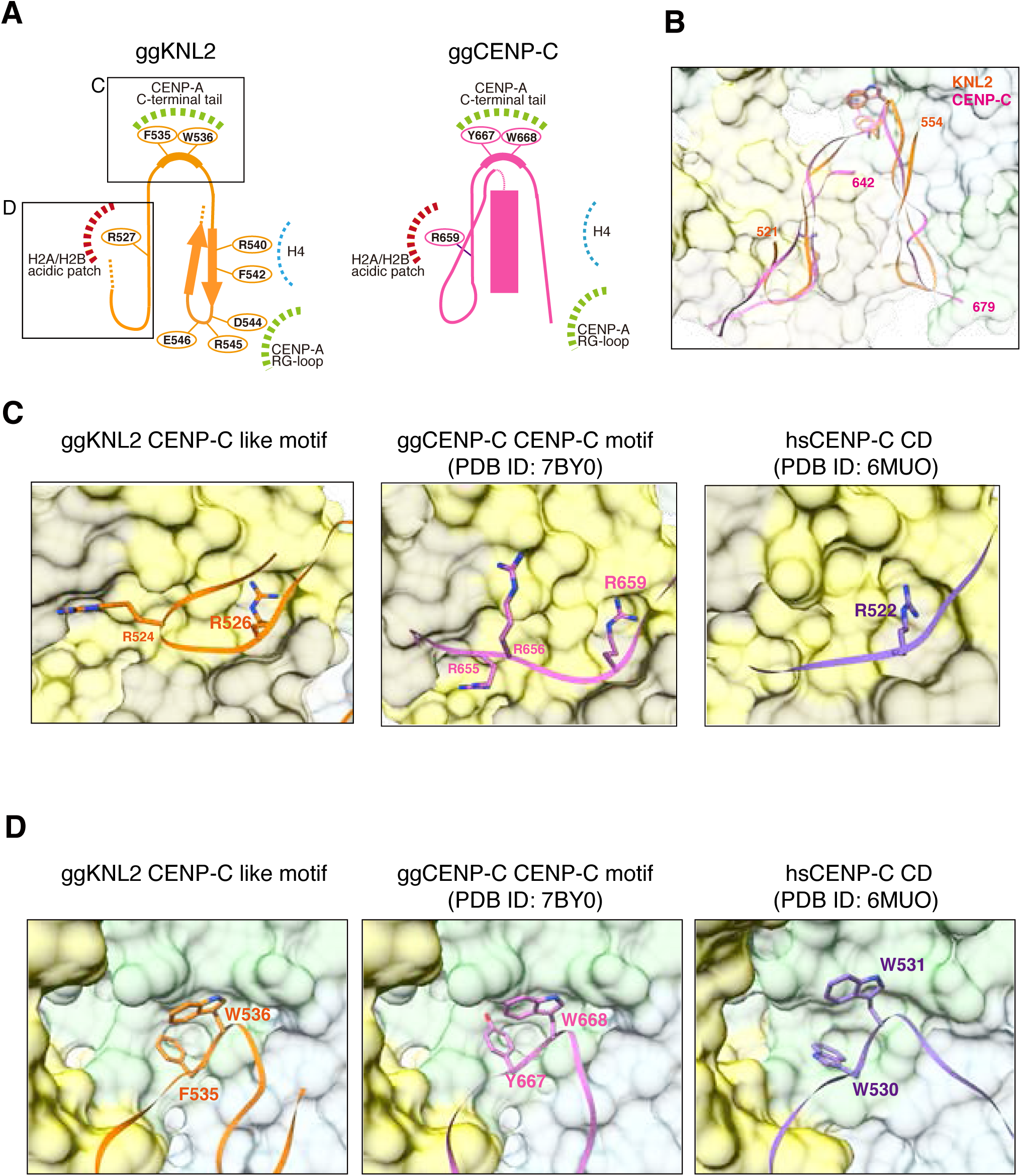
Structural comparison of the CENP-A nucleosome binding between KNL2 and CENP-C. A) Schematic representations of the structure of the CENP-A nucleosome binding region in chicken KNL2 (ggKNL2, orange) and chicken CENP-C (ggCENP-C, magenta). Detailed comparison of the H2A/H2B acidic patch and CENP-A C-terminal tail binding sites, indicated by squares, are presented in C and D. **B)** Structural comparison of the CENP-A nucleosome binding interface between KNL2 and CENP-C. The backbone structures of ggKNL2 (residues 521-554, orange) and ggCENP-C (residues 642-679, PDB ID: 7BY0, magenta) are superimposed. The CENP-A nucleosome is presented in a surface model. **C)** Structural comparison of the H2A/H2B acidic patch recognition mode among ggKNL2 CENP-C like motif, ggCENP-C CENP-C motif (PDB ID: 7BY0) and the human CENP-C central domain (CD) (PDB ID: 6MUO) on the CENP-A nucleosome. **D)** Structural comparison of the CENP-A C-terminal tail binding region among the ggKNL2 CENP-C like motif, ggCENP-C CENP-C motif and the human CENP-C CD on the CENP-A nucleosome.

**Figure EV4.**
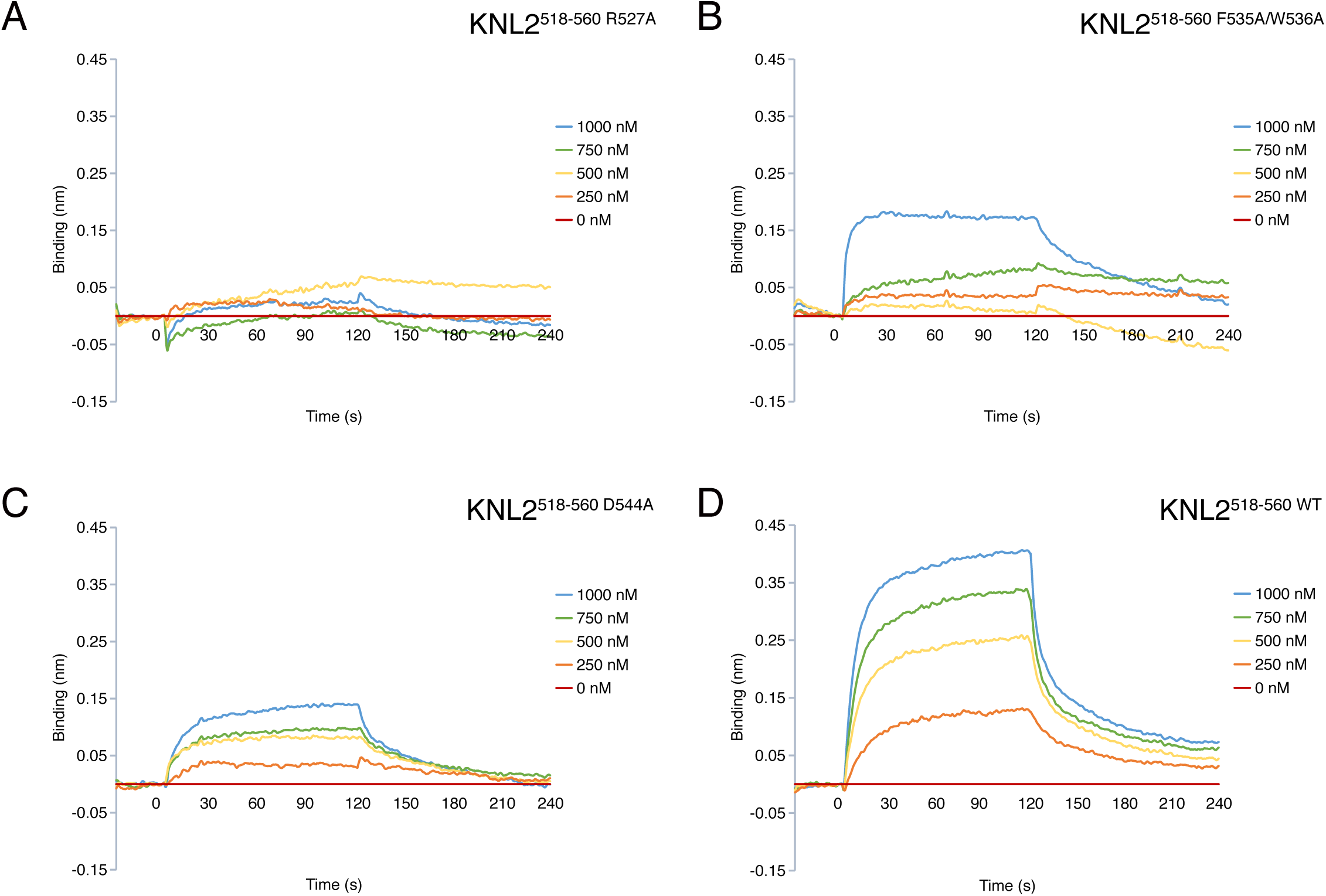
BLI analyses of the CENP-A nucleosome binding of KNL2^518-560^ mutants. BLI assays to examine the interaction between GST-KNL2^518-560^ mutants (immobilized ligand) and CENP-A nucleosome (analyte): **A)**, GST-KNL2^518-560^ R527A mutant; **B)**, GST- KNL2^518-560^ F535A/W536A; **C)**, GST-KNL2^518-560^ D544A; D) GST-KNL2^518-560^ wild type. Representative BLI sensorgrams measured with analyte concentrations ranging from 0 to 1 μM are plotted. The BLI sensorgrams were measured at association time intervals of 120 s and dissociation time intervals of 120 s.

## References

Ali-Ahmad A, Bilokapic S, Schafer IB, Halic M, Sekulic N (2019) CENP-C unwraps the human CENP-A nucleosome through the H2A C-terminal tail. EMBO reports 20: e48913

Allshire RC, Karpen GH (2008) Epigenetic regulation of centromeric chromatin: old dogs, new tricks? Nature reviews Genetics 9: 923–937

Allu PK, Dawicki-McKenna JM, Van Eeuwen T, Slavin M, Braitbard M, Xu C, Kalisman N, Murakami K, Black BE (2019) Structure of the Human Core Centromeric Nucleosome Complex. Current biology : CB 29: 2625–2639 e2625

Amano M, Suzuki A, Hori T, Backer C, Okawa K, Cheeseman IM, Fukagawa T (2009) The CENP-S complex is essential for the stable assembly of outer kinetochore structure. The Journal of cell biology 186: 173–182

Arimura Y, Tachiwana H, Oda T, Sato M, Kurumizaka H (2012) Structural analysis of the hexasome, lacking one histone H2A/H2B dimer from the conventional nucleosome. Biochemistry 51: 3302–3309

Arimura Y, Tachiwana H, Takagi H, Hori T, Kimura H, Fukagawa T, Kurumizaka H (2019) The CENP-A centromere targeting domain facilitates H4K20 monomethylation in the nucleosome by structural polymorphism. Nature communications 10: 576

Ariyoshi M, Makino F, Watanabe R, Nakagawa R, Kato T, Namba K, Arimura Y, Fujita R, Kurumizaka H, Okumura EI et al (2021) Cryo-EM structure of the CENP-A nucleosome in complex with phosphorylated CENP-C. The EMBO journal 40: e105671

Black BE, Cleveland DW (2011) Epigenetic centromere propagation and the nature of CENP-a nucleosomes. Cell 144: 471–479

Carroll CW, Silva MC, Godek KM, Jansen LE, Straight AF (2009) Centromere assembly requires the direct recognition of CENP-A nucleosomes by CENP-N. Nature cell biology 11: 896–902

Chittori S, Hong J, Saunders H, Feng H, Ghirlando R, Kelly AE, Bai Y, Subramaniam S (2018) Structural mechanisms of centromeric nucleosome recognition by the kinetochore protein CENP-N. Science 359: 339–343

Dambacher S, Deng W, Hahn M, Sadic D, Frohlich J, Nuber A, Hoischen C, Diekmann S, Leonhardt H, Schotta G (2012) CENP-C facilitates the recruitment of M18BP1 to centromeric chromatin. Nucleus 3: 101–110

Dunleavy EM, Roche D, Tagami H, Lacoste N, Ray-Gallet D, Nakamura Y, Daigo Y, Nakatani Y, Almouzni-Pettinotti G (2009) HJURP is a cell-cycle-dependent maintenance and deposition factor of CENP-A at centromeres. Cell 137: 485–497

Dyer PN, Edayathumangalam RS, White CL, Bao Y, Chakravarthy S, Muthurajan UM, Luger K (2004) Reconstitution of nucleosome core particles from recombinant histones and DNA. Methods in enzymology 375: 23–44

Emsley P, Lohkamp B, Scott WG, Cowtan K (2010) Features and development of Coot. Acta crystallographica Section D, Biological crystallography 66: 486–501

Falk SJ, Guo LY, Sekulic N, Smoak EM, Mani T, Logsdon GA, Gupta K, Jansen LE, Van Duyne GD, Vinogradov SA et al (2015) Chromosomes. CENP-C reshapes and stabilizes CENP-A nucleosomes at the centromere. Science 348: 699–703

Fang J, Liu Y, Wei Y, Deng W, Yu Z, Huang L, Teng Y, Yao T, You Q, Ruan H et al (2015) Structural transitions of centromeric chromatin regulate the cell cycle-dependent recruitment of CENP-N. Genes & development 29: 1058–1073

Foltz DR, Jansen LE, Bailey AO, Yates JR, 3rd, Bassett EA, Wood S, Black BE, Cleveland DW (2009) Centromere-specific assembly of CENP-a nucleosomes is mediated by HJURP. Cell 137: 472–484

Foltz DR, Jansen LE, Black BE, Bailey AO, Yates JR, 3rd, Cleveland DW (2006) The human CENP-A centromeric nucleosome-associated complex. Nature cell biology 8: 458–469

French BT, Straight AF (2019) CDK phosphorylation of Xenopus laevis M18BP1 promotes its metaphase centromere localization. The EMBO journal 38

French BT, Westhorpe FG, Limouse C, Straight AF (2017) Xenopus laevis M18BP1 Directly Binds Existing CENP-A Nucleosomes to Promote Centromeric Chromatin Assembly. Developmental cell 42: 190–199 e110

Fujita Y, Hayashi T, Kiyomitsu T, Toyoda Y, Kokubu A, Obuse C, Yanagida M (2007) Priming of centromere for CENP-A recruitment by human hMis18alpha, hMis18beta, and M18BP1. Developmental cell 12: 17–30

Fukagawa T, Earnshaw WC (2014) The Centromere: Chromatin Foundation for the Kinetochore Machinery. Developmental cell 30: 496–508

Hayashi T, Fujita Y, Iwasaki O, Adachi Y, Takahashi K, Yanagida M (2004) Mis16 and Mis18 are required for CENP-A loading and histone deacetylation at centromeres. Cell 118: 715–729

Hori T, Amano M, Suzuki A, Backer CB, Welburn JP, Dong Y, McEwen BF, Shang WH, Suzuki E, Okawa K et al (2008) CCAN makes multiple contacts with centromeric DNA to provide distinct pathways to the outer kinetochore. Cell 135: 1039–1052

Hori T, Cao J, Nishimura K, Ariyoshi M, Arimura Y, Kurumizaka H, Fukagawa T (2020) Essentiality of CENP-A Depends on Its Binding Mode to HJURP. Cell reports 33: 108388

Hori T, Shang WH, Hara M, Ariyoshi M, Arimura Y, Fujita R, Kurumizaka H, Fukagawa T (2017) Association of M18BP1/KNL2 with CENP-A Nucleosome Is Essential for Centromere Formation in Non-mammalian Vertebrates. Developmental cell 42: 181–189 e183

Izuta H, Ikeno M, Suzuki N, Tomonaga T, Nozaki N, Obuse C, Kisu Y, Goshima N, Nomura F, Nomura N et al (2006) Comprehensive analysis of the ICEN (Interphase Centromere Complex) components enriched in the CENP-A chromatin of human cells. Genes to cells : devoted to molecular & cellular mechanisms 11: 673–684

Jansen LE, Black BE, Foltz DR, Cleveland DW (2007) Propagation of centromeric chromatin requires exit from mitosis. The Journal of cell biology 176: 795–805

Kato H, Jiang J, Zhou BR, Rozendaal M, Feng H, Ghirlando R, Xiao TS, Straight AF, Bai Y (2013a) A conserved mechanism for centromeric nucleosome recognition by centromere protein CENP-C. Science 340: 1110–1113

Kato H, Zhou BR, Feng H, Bai Y (2013b) An evolving tail of centromere histone variant CENP-A. Cell cycle 12: 3133–3134

Klare K, Weir JR, Basilico F, Zimniak T, Massimiliano L, Ludwigs N, Herzog F, Musacchio A (2015) CENP-C is a blueprint for constitutive centromere-associated network assembly within human kinetochores. The Journal of cell biology 210: 11–22

Kral L (2015) Possible identification of CENP-C in fish and the presence of the CENP-C motif in M18BP1 of vertebrates. F1000Res 4: 474

Liebschner D, Afonine PV, Baker ML, Bunkoczi G, Chen VB, Croll TI, Hintze B, Hung LW, Jain S, McCoy AJ et al (2019) Macromolecular structure determination using X-rays, neutrons and electrons: recent developments in Phenix. Acta Crystallogr D Struct Biol 75: 861–877

Maddox PS, Hyndman F, Monen J, Oegema K, Desai A (2007) Functional genomics identifies a Myb domain-containing protein family required for assembly of CENP-A chromatin. The Journal of cell biology 176: 757–763

Mastronarde DN (2005) Automated electron microscope tomography using robust prediction of specimen movements. Journal of structural biology 152: 36–51

McKinley KL, Cheeseman IM (2014) Polo-like Kinase 1 Licenses CENP-A Deposition at Centromeres. Cell 158: 397–411

Moree B, Meyer CB, Fuller CJ, Straight AF (2011) CENP-C recruits M18BP1 to centromeres to promote CENP-A chromatin assembly. The Journal of cell biology 194: 855–871

Nardi IK, Zasadzinska E, Stellfox ME, Knippler CM, Foltz DR (2016) Licensing of Centromeric Chromatin Assembly through the Mis18alpha-Mis18beta Heterotetramer. Molecular cell 61: 774–787

Nishino T, Takeuchi K, Gascoigne KE, Suzuki A, Hori T, Oyama T, Morikawa K, Cheeseman IM, Fukagawa T (2012) CENP-T-W-S-X forms a unique centromeric chromatin structure with a histone-like fold. Cell 148: 487–501

Okada M, Cheeseman IM, Hori T, Okawa K, McLeod IX, Yates JR, 3rd, Desai A, Fukagawa T (2006) The CENP-H-I complex is required for the efficient incorporation of newly synthesized CENP-A into centromeres. Nature cell biology 8: 446–457

Pan D, Klare K, Petrovic A, Take A, Walstein K, Singh P, Rondelet A, Bird AW, Musacchio A (2017) CDK-regulated dimerization of M18BP1 on a Mis18 hexamer is necessary for CENP-A loading. eLife 6

Pan D, Walstein K, Take A, Bier D, Kaiser N, Musacchio A (2019) Mechanism of centromere recruitment of the CENP-A chaperone HJURP and its implications for centromere licensing. Nature communications 10: 4046

Pentakota S, Zhou K, Smith C, Maffini S, Petrovic A, Morgan GP, Weir JR, Vetter IR, Musacchio A, Luger K (2017) Decoding the centromeric nucleosome through CENP-N. eLife 6

Perpelescu M, Fukagawa T (2011) The ABCs of CENPs. Chromosoma 120: 425–446

Perpelescu M, Hori T, Toyoda A, Misu S, Monma N, Ikeo K, Obuse C, Fujiyama A, Fukagawa T (2015) HJURP is involved in the expansion of centromeric chromatin. Molecular biology of the cell 26: 2742–2754

Pesenti ME, Raisch T, Conti D, Walstein K, Hoffmann I, Vogt D, Prumbaum D, Vetter IR, Raunser S, Musacchio A (2022) Structure of the human inner kinetochore CCAN complex and its significance for human centromere organization. Molecular cell

Pettersen EF, Goddard TD, Huang CC, Couch GS, Greenblatt DM, Meng EC, Ferrin TE (2004) UCSF Chimera--a visualization system for exploratory research and analysis. J Comput Chem 25: 1605–1612

Sandmann M, Talbert P, Demidov D, Kuhlmann M, Rutten T, Conrad U, Lermontova I (2017) Targeting of A. thaliana KNL2 to centromeres depends on the conserved CENPC-k motif in its C-terminus. The Plant cell

Silva MC, Bodor DL, Stellfox ME, Martins NM, Hochegger H, Foltz DR, Jansen LE (2012) Cdk activity couples epigenetic centromere inheritance to cell cycle progression. Developmental cell 22: 52–63

Spiller F, Medina-Pritchard B, Abad MA, Wear MA, Molina O, Earnshaw WC, Jeyaprakash AA (2017) Molecular basis for Cdk1-regulated timing of Mis18 complex assembly and CENP-A deposition. EMBO reports 18: 894–905

Stellfox ME, Nardi IK, Knippler CM, Foltz DR (2016) Differential Binding Partners of the Mis18alpha/beta YIPPEE Domains Regulate Mis18 Complex Recruitment to Centromeres. Cell reports 15: 2127–2135

Subramanian L, Medina-Pritchard B, Barton R, Spiller F, Kulasegaran-Shylini R, Radaviciute G, Allshire RC, Arockia Jeyaprakash A (2016) Centromere localization and function of Mis18 requires Yippee-like domain-mediated oligomerization. EMBO reports

Tachiwana H, Kagawa W, Shiga T, Osakabe A, Miya Y, Saito K, Hayashi-Takanaka Y, Oda T, Sato M, Park SY et al (2011) Crystal structure of the human centromeric nucleosome containing CENP-A. Nature 476: 232–235

Tanaka Y, Tawaramoto-Sasanuma M, Kawaguchi S, Ohta T, Yoda K, Kurumizaka H, Yokoyama S (2004) Expression and purification of recombinant human histones. Methods 33: 3–11

Tian T, Li X, Liu Y, Wang C, Liu X, Bi G, Zhang X, Yao X, Zhou ZH, Zang J (2018) Molecular basis for CENP-N recognition of CENP-A nucleosome on the human kinetochore. Cell research 28: 374–378

Watanabe R, Hara M, Okumura EI, Herve S, Fachinetti D, Ariyoshi M, Fukagawa T (2019) CDK1-mediated CENP-C phosphorylation modulates CENP-A binding and mitotic kinetochore localization. The Journal of cell biology 218: 4042–4062

Watanabe R, Hirano Y, Hara M, Hiraoka Y, Fukagawa T (2022) Mobility of kinetochore proteins measured by FRAP analysis in living cells. Chromosome research : an international journal on the molecular, supramolecular and evolutionary aspects of chromosome biology 30: 43–57

Weir JR, Faesen AC, Klare K, Petrovic A, Basilico F, Fischbock J, Pentakota S, Keller J, Pesenti ME, Pan D et al (2016) Insights from biochemical reconstitution into the architecture of human kinetochores. Nature 537: 249–253

Westhorpe FG, Fuller CJ, Straight AF (2015) A cell-free CENP-A assembly system defines the chromatin requirements for centromere maintenance. The Journal of cell biology 209: 789–801

Westhorpe FG, Straight AF (2013) Functions of the centromere and kinetochore in chromosome segregation. Current opinion in cell biology 25: 334–340

Yan K, Yang J, Zhang Z, McLaughlin SH, Chang L, Fasci D, Ehrenhofer-Murray AE, Heck AJR, Barford D (2019) Structure of the inner kinetochore CCAN complex assembled onto a centromeric nucleosome. Nature 574: 278–282

Yatskevich S, Muir KW, Bellini D, Zhang Z, Yang J, Tischer T, Predin M, Dendooven T, McLaughlin SH, Barford D (2022) Structure of the human inner kinetochore bound to a centromeric CENP-A nucleosome. Science: eabn3810

Zhang K (2016) Gctf: Real-time CTF determination and correction. Journal of structural biology 193: 1–12

Zheng SQ, Palovcak E, Armache JP, Verba KA, Cheng Y, Agard DA (2017) MotionCor2: anisotropic correction of beam-induced motion for improved cryo-electron microscopy. Nature methods 14: 331–332

Zivanov J, Nakane T, Forsberg BO, Kimanius D, Hagen WJ, Lindahl E, Scheres SH (2018) New tools for automated high-resolution cryo-EM structure determination in RELION-3. eLife 7

